# Expression of a human cDNA in moss results in spliced mRNAs and fragmentary protein isoforms

**DOI:** 10.1101/2020.09.30.320721

**Authors:** Oguz Top, Stella W. L. Milferstaedt, Nico van Gessel, Sebastian N. W. Hoernstein, Bugra Özdemir, Eva L. Decker, Ralf Reski

## Abstract

Production of biopharmaceuticals relies on the expression of mammalian cDNAs in host organisms. Here we show that the expression of a human cDNA in the moss *Physcomitrella patens* generates the expected full-length and four additional transcripts due to unexpected splicing. This mRNA splicing results in non-functional protein isoforms, cellular misallocation of the proteins and low product yields. We integrated these results together with the results of our analysis of all 32,926 protein-encoding *P. patens* genes and their 87,533 annotated transcripts in a web application, physCO, for automatized codon-optimization. A thus optimized cDNA results in about eleven times more protein, which correctly localizes to the ER. An analysis of codon preferences of different production hosts suggests that similar effects also occur in non-plant hosts. We anticipate that the use of our methodology will prevent so far undetected mRNA heterosplicing resulting in maximized functional protein amounts for basic biology and biotechnology.

## Introduction

The majority of eukaryotic genes comprise multiple exons interspersed by noncoding introns, the number and length of which vary between species. The human genome size (GRCh38.p13, Genome Reference Consortium Human Build 38 patch release 13) is 3,099,706,400 bp compromising 20,444 protein-encoding genes, with 8.8 exons and 7.8 introns per gene on average^1^. Generally, human genes have many short exons (average 50 codons) separated by long introns (up to >10 kbp)^2^. It is widely accepted that such an exon-intron gene structure emerged during evolution and lead to complex gene regulation and diversification of multicellular eukaryotes^2,3^. Introns that may contain regulatory elements affecting gene expression are removed from pre-mRNAs by RNA splicing. Subsequently, exons are ligated together to generate functional mature mRNAs^4^. Over 30 years of research on pre-mRNA splicing elucidated an underlying complex regulation, which includes exon versus intron recognition^5^, co-transcriptional splicing^6^, alternative splicing^7^, exonic and intronic splicing enhancers and repressors^8,9^, and mRNA export from the nucleus^10^.

Alternative splicing (AS) is a fundamental regulatory process that contributes to proteome expansion^11^. AS can produce multiple mRNAs from the same gene through the preference of splice sites during pre-mRNA splicing. It is modulated based on sequence motifs in the pre-mRNA, the interactions between RNA-binding proteins and splice sites, different cell types, and environmental signals^12^. Splicing is carried out by a large ribonucleoprotein (RNP) complex, the spliceosome, which brings selected exons together by two transesterification reactions^13^. During splicing, uridine (U)-rich small nuclear RNPs (snRNPs) together with non-snRNP splicing factors, and serine/arginine-rich (SR) proteins participate in the recognition of 5’- and 3’-splice sites as well as the branch site^14^. The core of the spliceosome is highly conserved in all well-characterized eukaryotes^3,15^. The key requirement for splicing is the recognition of a donor splice site by the U1 snRNP^16^. On the other hand, the acceptor splice site, downstream of the polypyrimidine tract, is recognized by the U5 snRNP^17^. Another important splicing element containing the reactive adenosine, the branch point signal, is located upstream of the acceptor splice signal and is important for the formation of the lariat-like intermediate structure^17^. While the majority of the introns have “GT” at the donor splice site and “AG” at the acceptor splice site, in rare cases introns can have unusual dinucleotides: for instance “AT” at the donor site and “AC” at the acceptor site^18^. This class of introns is spliced out of the pre-mRNA by the minor spliceosome U12^19^.

Besides its role in proteome expansion, AS controls transcript levels by generating unstable mRNA isoforms that may activate nonsense-mediated mRNA decay (NMD)^20^. Generation of alternative functional mRNAs due to the introduction of premature termination codons (PTC), intron retention, and alternative use of the 5’- and 3’-splice sites causes changes in the localization, stability and/or function of a protein. While in humans more than 95% of genes encode alternatively spliced mRNAs^21^, in an early study 6,556 out of 39,106 predicted genes showed AS in the basic land plant *Physcomitrella patens* (moss) with a strong bias towards intron retention^22^. Many PTC-containing alternatively spliced transcripts are NMD targets in *P. patens*^23^. Furthermore, about half of the AS events altered coding sequences and resulted in 2,380 distinct proteins. In these studies, analysis of AS relied on expressed sequence tags (ESTs) and the alignment of ESTs and cDNA sequences to the genome. The percentage of AS events in *P. patens* is estimated as 58%, but large-scale and/or tissue-dependent profiling of AS in moss by the use of improved sequencing methods may result in a higher percentage^24^.

Two models of splice-site recognition exist^25^: In the intron-definition model, U1 binds to the upstream donor splice site and U2AF/U2 to the downstream acceptor splice site and branch site of the same intron^26,27^. The length of introns, which probably remain short under evolutionary selection^3^, limits the efficiency of splicing. In lower eukaryotes where introns are shorter, this mechanism might be dominant^15^. In the exon-definition model, the splicing machinery recognizes splice sites flanking the same exon^28,29^. This is the major mechanism in higher eukaryotes, where introns are generally longer. Due to these differences, the correct splicing of transgene transcripts has to be considered to maximize the production of a functional protein. For instance, it was necessary to remove a cryptic intron in the coding sequence (CDS) of green fluorescent protein (GFP) from the jellyfish *Aequorea victoria* to function in the flowering plant *Arabidopsis thaliana*^30^.

The use of plants as alternative pharmaceutical protein production hosts has been on the rise for the past two decades, and some plant-made products have reached the market^31^. However, the percentage of plant-based biopharmaceuticals in the pharmaceutical market is still minute, mainly because of lower yields compared to prokaryotic and mammalian systems^32^. Nevertheless, they offer alternative solutions in niche areas^33^. Production of blood clotting factors could be such a niche^31^. The human blood-clotting factor IX (FIX), a member of the intrinsic pathway of coagulation, is required for normal hemostasis and its absence or abnormal levels causes hemophilia B^34^. Current treatment is restricted to protein replacement therapy and factor concentrates costs are between $100,000 and $200,000 per patient and year^35^. Thus, there is a need for an economic production of FIX, which makes us attempting to produce it in the moss system. The moss *P. patens,* a model organism for evolutionary and functional genomics approaches, is an established host for the production of complex recombinant biopharmaceuticals and proven its suitability for large-scale production^36,37^. The first moss-made drug candidate, moss-aGal for enzyme replacement therapy of Fabry disease, has successfully completed phase I clinical trials^38^. Additionally, a variety of other potential biopharmaceuticals is being produced in moss^39–41^.

In the present study, we report the expression of a human blood clotting factor IX-encoding cDNA in *P. patens*. We show that specific sequence motifs in the FIX mRNA were recognized by the splicing machinery as donor and acceptor sites and produced several different transcripts, which resulted in different protein isoforms. We named this new phenomenon heterosplicing, and were able to prevent it by changing the splice sites and optimizing codons without affecting the FIX aa sequence (optiFIX). Expression of optiFIX resulted in full-length transcripts only, and an eleven-fold increase of protein accumulation. In addition, we found heterosplicing also in transgenic moss lines expressing the CDS of human factor H (FH) CDS. By optimizing the sequence, we generated optiFH, which showed no heterosplicing and yielded significantly enhanced FH protein abundances *in vivo.* We suggest that our methodology of CDS optimization will provide a better quantification of transcript levels, and precise detection, purification, and quantification of functional proteins. It may reduce the bioenergetic costs of heterologous cDNA expression by preventing the formation of non-functional products and maximizing functional recombinant protein production, major bottlenecks in plant-based systems.

## RESULTS

### Predicted miRNA activity is inhibited by mutating miRNA binding sites

MicroRNAs can regulate gene expression at the posttranscriptional level, either via target mRNA degradation or translational inhibition. The possible actions of *P. patens* miRNAs on the RNA sequence for human Factor IX (NCBI reference NM_000133) were investigated using psRNATarget (http://plantgrn.noble.org/v1_psRNATarget/). Thus, two miRNAs were identified (miR1028c-3p and miR1044-3p), that might interfere with FIX expression in this moss. The respective binding sites on the FIX RNA sequences were modified in the DNA construct without a change in the FIX aa sequence. The potential binding site of ppt-miR1028c-3p, which might inhibit translation, was changed into ATCATGTGAGCCAGCTGTACCC between positions 495 and 516. The binding site of ppt-miR1044-3p, which might lead to cleavage of the FIX RNA, was mutated into AGGGTAAGTACGGCATCTAT between positions 1304 and 1323. The sequence of miRNAs and their binding sites as well as the maximum energy to unpair the target site (UPE) and expectation value are shown in Supplementary Table 1.

### Splicing of FIX mRNA occurs in transgenic plants and in transiently transfected cells

The native FIX signal peptide was replaced by the moss aspartic proteinase signal peptide of *Pp*AP1 to target the protein to the secretory pathway for its proper posttranslational modifications^42^. After transfection of the parental moss line Δ*xt/ft*^41^ with the human FIX-encoding, cDNA-based expression construct that comprises the complete FIX coding sequence (CDS) of 1401 bp without any introns; 49 transgenic moss lines were regenerated and screened. The presence and integration of the full-length FIX CDS in the moss genome was confirmed via PCR (Supplementary Fig. 1). In order to validate the completeness of the FIX transcripts, RT-PCR was performed with primers spanning the complete CDS. In addition to the full-length transcript, several smaller products occurred (Fig. 1). In cDNA from the juvenile moss protonemal tissue, PCR products of discrete sizes were detected ranging from ~700 to 1370 bp (Fig. 1a). Sequencing of the PCR products revealed that five of the observed bands were FIX-specific products with a length of 1371 bp, 1240 bp, 895 bp, 769 bp, and 691 bp, respectively. Apparently, unexpected splicing of the FIX mRNA produced the smaller aberrant transcripts. To analyze this phenomenon across different tissues, we repeated the same procedure with RNA isolated from adult plants (gametophores). FIX transcripts detected in gametophores were consistent with those from protonema (Fig. 1b).

**Fig. 1:**
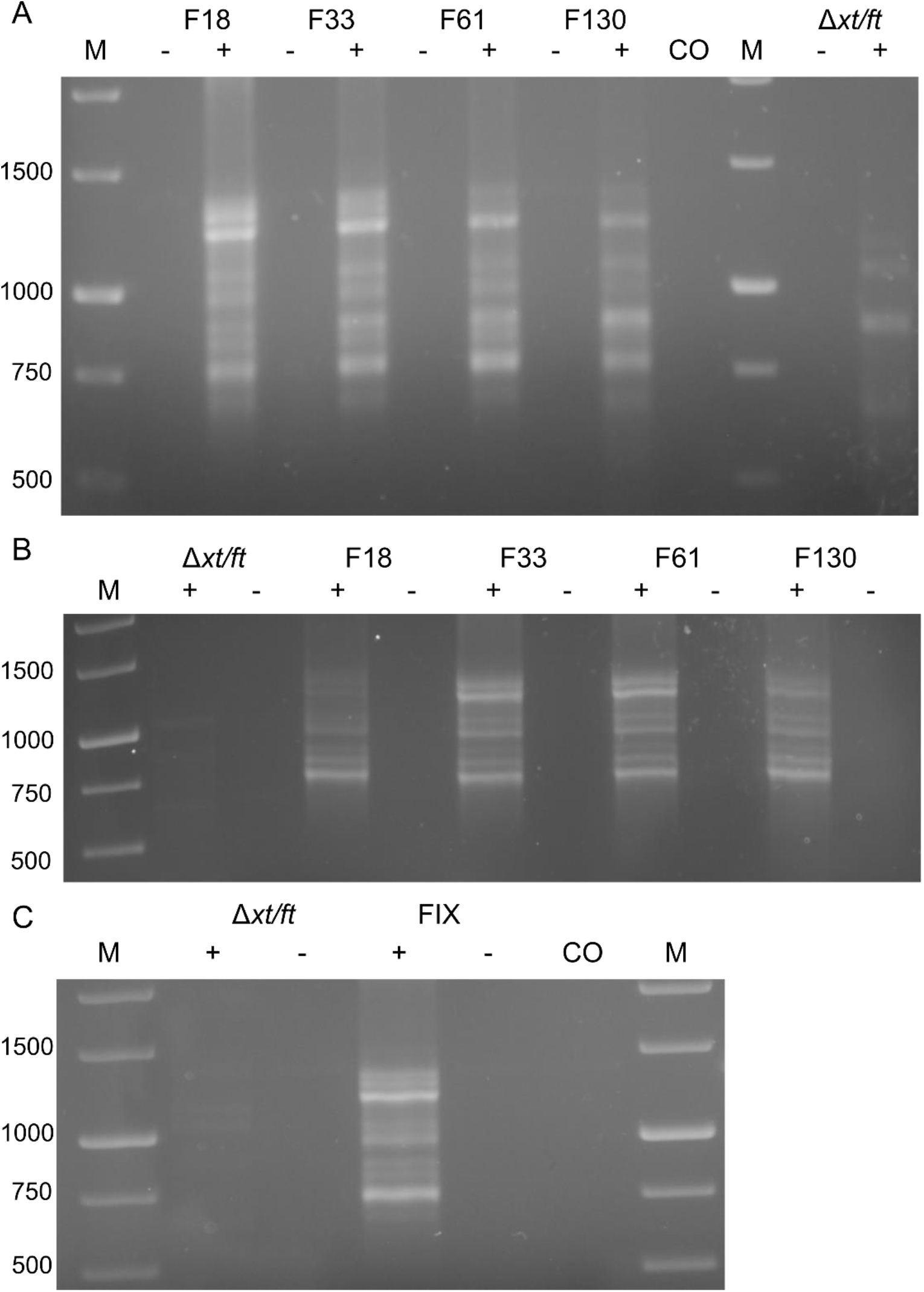
Expression of FIX in *Physcomitrella patens*. **a** Total RNA from protonema cultures of 4 FIX-transgenic lines (F) and the parental line Δ*xt/ft* was prepared and used for RT-PCR using the primers FIXfwdB and FIXrevB. **b** RT-PCR of FIX using the primers FIXfwdB and FIXrevB from gametophores of the same lines. **c** RT-PCR of FIX with mRNA extracted from transiently transfected (FIX) as well as non-transfected (Δ*xt/ft*) cells. M: 1 kb Marker (Thermo Fisher Scientific), -: without reverse transcription, +: with reverse transcription, CO: Water control.

The sequencing of different PCR products revealed that specific motifs along the human FIX mRNA were recognized as splice signals in *P. patens.* The spliced transcripts were identical in all 49 transgenic lines and in 2 different moss tissues. Hence, we conclude that different transcripts were exclusively derived from unexpected splicing of the mRNA transcribed from the complete FIX CDS, and not by a partial integration of the FIX expression construct into the *P. patens* genome. PCRs from genomic DNA support this conclusion (Supplementary Fig. 1).

In order to scrutinize the presence of spliced FIX CDS variants in transiently transfected cells (protoplasts), total RNA was extracted from cells 14 days after transfection. After RT-PCR, FIX CDS variants with varying lengths occurred. Like in the RT-PCR from transgenic lines, 5 FIX CDS variants (1 full-length and 4 aberrant spliced FIX transcripts) were confirmed in these protoplasts (Fig. 1c). Thus, we found an unexpected splicing of an mRNA transcribed from a human full-length cDNA in moss, a phenomenon we henceforth name heterosplicing.

### Spliced FIX transcripts and splice-site analysis

Splice junctions within the FIX mRNA were revealed by sequencing. The splicing of the primary transcripts from human FIX cDNA in moss caused the production of four alternative products (Fig. 2a): i. deletion of a 44 aa long domain in the part coding for the heavy chain (Fig. 2a.ii), ii. large deletion in the part coding for the light chain (115 aa) together with the absence observed in the heavy chain (Fig. 2a.iii), iii. nearly complete absence of the sequence coding for the light chain and parts of the activation peptide domain (157 aa) and the heavy chain (Fig. 2a.iv), iv. nearly complete lack of the sequence encoding the light chain, complete absence of the activation peptide domain (186 aa) and the sequence representing 41 aa of the heavy chain (Fig. 2a.v). The heavy chain harbors a serine protease domain, which is responsible for the activation of Factor X in the presence of Factor VIII, calcium and phospholipid surfaces following removal of the activation peptide. Any deletion in this domain will interfere with the activity of the FIX protein. The light chain consists of the Gla domain followed by two EGF domains. Any loss in these domains will adversely affect the properties of FIX. Therefore, any alterations caused by the heterosplicing will result in the loss of FIX function.

**Fig. 2:**
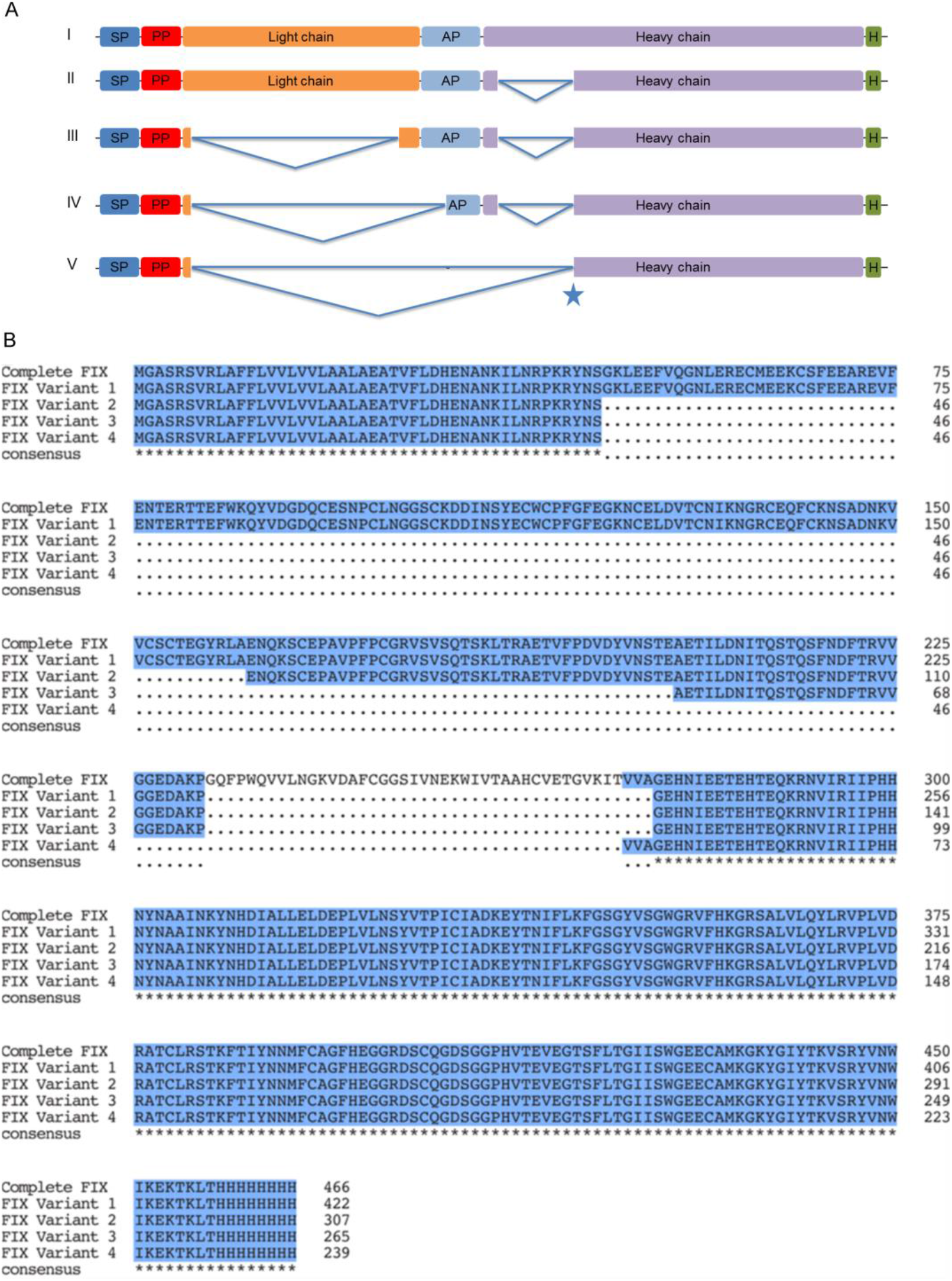
Schematic representation of FIX transcripts and multiple sequence alignment of predicted FIX protein isoforms based on experimentally verified transcripts detected in *Physcomitrella patens*. **a** Full-length FIX transcript (a.i), FIX variant with the deletion of a 44 aa long domain in the CDS for the heavy chain (a.ii), FIX variant with deletions in the CDS for both light and heavy chain (a.iii), FIX variant with deletions in the CDS for light chain, activation peptide and heavy chain (a.iv), FIX variant missing the CDS completely for the activation peptide, nearly full-length of light chain and 41 aa long domain in the heavy chain (a.v). The star indicates the usage of a donor site 9 bp earlier than previously identified donor site in the heavy chain. SP: PpAP1 signal peptide, PP: propeptide, AP: activation peptide, H: 8x His tag. **b** Alignment was performed and formatted for publication with MUSCLE via the “MSA” package for R^49,50^.

The GT-AG splicing rule applies for almost all eukaryotic genes^43^ and especially *P. patens* accepts control elements originally optimized for mammalian expression systems without a need for adapting these elements^44^. Here, we inferred four different motifs for acceptor and donor sites based on sequencing data. All FIX heterosplice sites followed the GT-AG rule. Moreover, the data revealed that a CAGGT motif is present in the exon-intron junction, i.e. the donor site (Fig. 3a). In the intron-exon junction, the acceptor site, the motif CAG is conserved in 3 out of 4 cases, but the adjacent two nucleotides do not follow any trend.

**Fig. 3:**
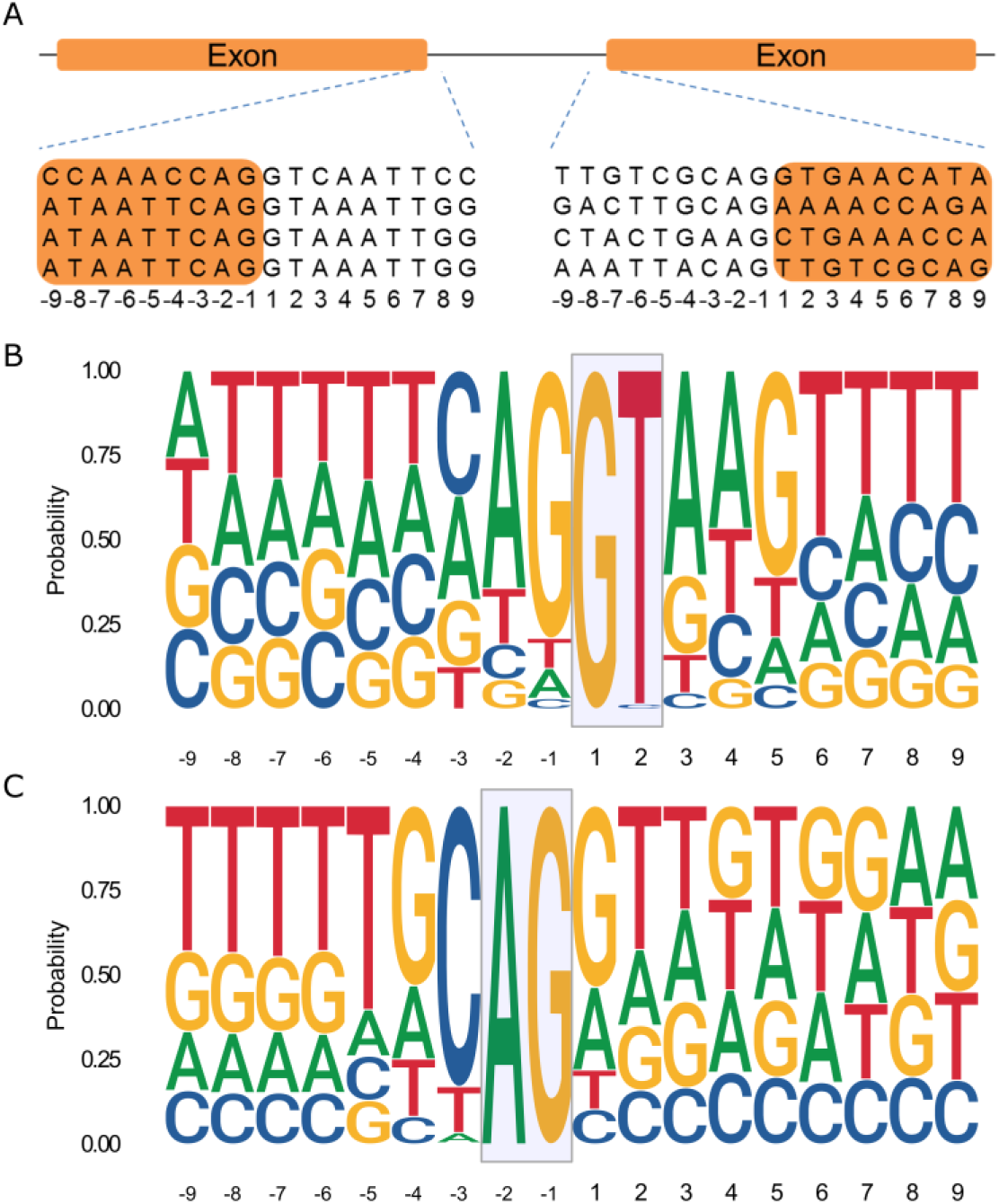
Schematic representation of donor and acceptor sites within the FIX sequence and consensus sequence motifs of donor and acceptor splice sites recovered from all 87,533 *P. patens* transcripts. **a** Donor and acceptor sites. Four different splicing motifs were identified after sequencing FIX variants. Left side shows donor site and right side shows acceptor site. “-1” represents last nucleotide of exon in the donor site (left side) or last nucleotide of intron in the acceptor site (right side). **b, c** Consensus genomic sequence motifs of donor (b) and acceptor splice sites (c) extracted from all 32,926 *P. patens* protein-coding genes. Probability and height of individual letters correspond to base frequencies at each position.

To analyze whether donor and acceptor sites and their neighboring nucleotides are conserved in *P. patens*, the genomic vicinity of all splice sites of all 87,533 annotated transcripts corresponding to the 32,926 protein-encoding genes of the current *P. patens* genome release v3.3^45^ were retrieved (Fig. *3*b, c). In alignment with our experimental findings, an *in silico*-analysis of these transcripts showed that the CAG|GT motif is the most abundant one in both donor and acceptor sites with a fraction of 23% and 18%, respectively. The full list of splice cites is compiled in Supplementary Tables 2 and 3.

Subsequently, we analyzed the FIX CDS with a moss splice-site prediction tool^46^, but it largely failed to predict the experimentally verified splicing motifs. This might be due to the fact that this tool has been developed several years ago using only 368 donor and acceptor sites^46^ and has not been updated since then.

### Detection of truncated mossFIX protein variants and validation by mass spectrometry

The analyses at RNA level suggested that *P. patens* would produce a full-length FIX protein and four aberrant fragmentary FIX isoforms. To validate this inference, the culture supernatants of transiently transfected cells were precipitated and analyzed with a polyclonal anti-FIX antibody via immunoblot after reduction and alkylation (Fig. 4a). Full-length FIX was expected at a molecular mass of around 52 kDa and spliced FIX variants were calculated based on the heterospliced transcripts with molecular masses of 47, 34, 29, and 27 kDa, respectively. Observed bands on the immunoblot could be attributed to these isoforms together with a band at around 40 kDa (potentially a degradation product) and a band at around 70 kDa (potentially dimers of fragmentary protein isoforms). Additionally, we used the other half of the supernatants for mass-spectrometric (MS) determination of FIX-variant peptide sequences. For the MS-database search, protein models for FIX variants derived from the sequenced splice variants were employed.

**Fig. 4:**
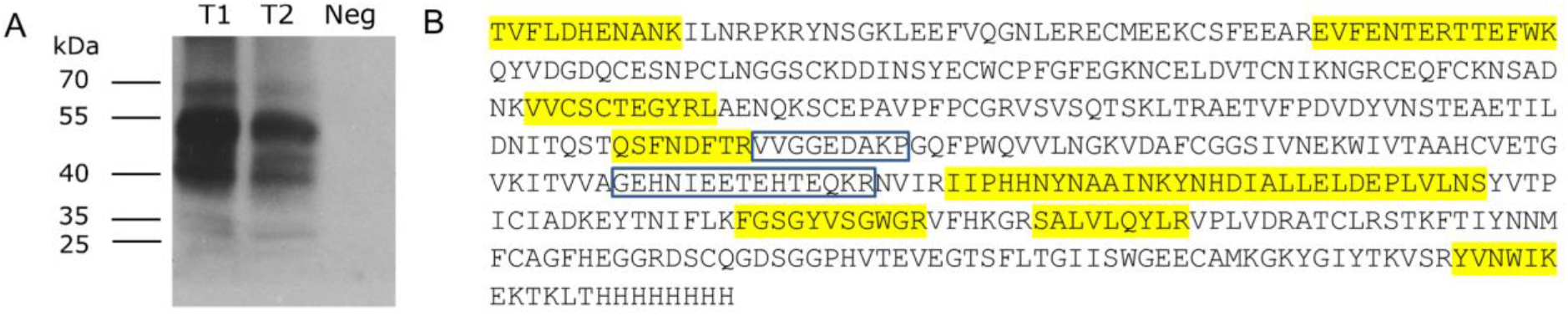
Analysis on moss-produced FIX protein isoforms. **a** Immunodetection of extracellular FIX produced by two transfections (anti-FIX Ab 1:5,000). Culture supernatant of non-transfected cells was used as negative control (Neg). T1, T2: Two different transient transfections with the FIX cDNA-based expression plasmid. **b** Full-length mossFIX amino acid sequence. Amino acids marked in yellow were identified in the MS analysis. Blue boxes mark the parts of the peptide VVGGEDAKPGEHNIEETEHTEQKR which was identified in MS analysis and represents a heterosplice product.

We detected the peptide VVGGEDAKPGEHNIEETEHTEQKR (Supplementary Fig. 2) in MS analysis after trypsin digestion, which resulted from splicing in the CDS for the heavy chain that led to the loss of 44 aa between the peptides VVGGEDAKP and GEHNIEETEHTEQKR (Fig. 4b; variants II, III, and IV in Fig. 2a). This result confirmed the presence of a predicted FIX variant caused by heterosplicing on an additional level.

### Prevention of heterosplicing by codon optimization

To analyze the effectiveness of mutating donor and acceptor sites on heterosplicing of the FIX mRNA, we created two different FIX CDS versions. In the first one, aspFIX, the codons neighboring the detected splice junctions were modified. *P. patens* was transiently transfected with the resulting aspFIX plasmid. On day 14, the cells were collected and RNA was isolated. RT-PCR and sequencing of the PCR products revealed that there were only two bands representing FIX variants: the full-length FIX CDS and a shorter variant (~700 bp) (Fig. 5a). The donor and acceptor sites of the shorter variant were identified (Fig. 5b) and this shorter variant was lacking the nucleotides encoding the light chain, AP and parts of the heavy chain (Fig. 5c). Moreover, splicing would cause a frameshift mutation resulting in early stop codons. Although such nonsense mRNAs may be degraded via nonsense-mediated mRNA decay (NMD), not all nonsense mRNAs undergo NMD, which was reported e.g. for *P. patens*^23^ and *A. thaliana*^47,48^. In the second attempt, we performed a complete codon optimization of the FIX CDS in the aspFIX construct according to previous studies^49,50^ resulting in the optiFIX sequence with an overall increased GC content. This was used to transiently transfect moss cells. Subsequent RT-PCR analysis and sequencing revealed that the full-length CDS of FIX was the only specific band; formation of any heterospliced FIX variant was not detected (Fig. 5d). The presence of exclusively the full-length CDS was confirmed on protein level via immunoblot analysis (Fig. 5e).

**Fig. 5:**
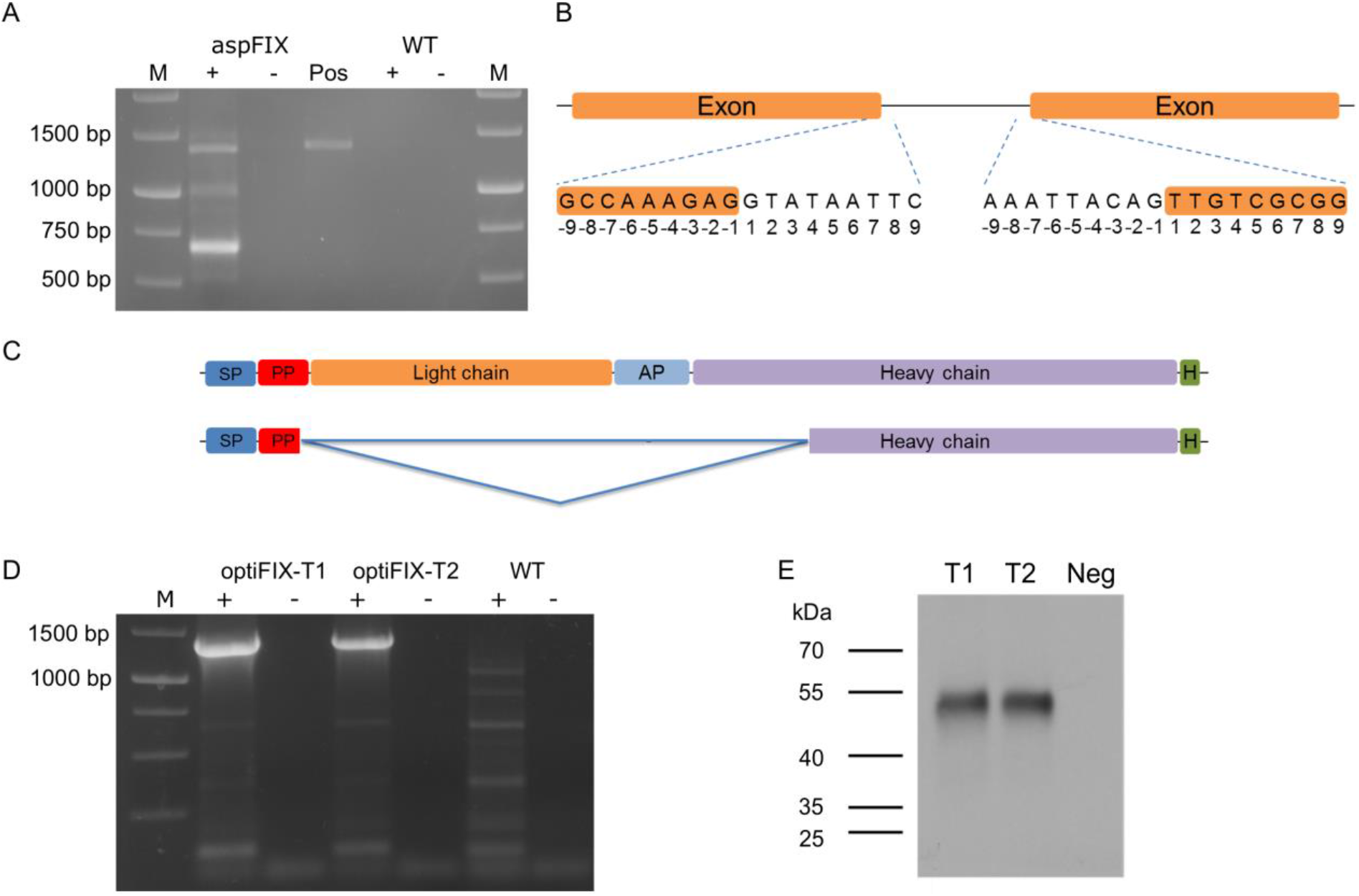
Heterosplicing of the human FIX CDS in *Physcomitrella patens* was successfully prevented by codon optimization. **a** RT-PCR of FIX RNA from cells that were transiently transfected with aspFIX and non-transfected cells (WT). M: 1 kb Marker (Thermo Fisher Scientific), −: without reverse transcription, +: with reverse transcription, Pos: PCR with aspFIX plasmid used as positive control. **b** Splicing motifs identified after sequencing the FIX variant. The left side shows the donor site and the right side the acceptor site. “−1” represents the last nucleotide of the exon at the donor site or the intron at the acceptor site, respectively. **c** Schematic representation of complete FIX (upper) and the additional variant (lower). SP: signal peptide, PP: propeptide, AP: activation peptide. **d** RT-PCR of FIX RNA from cells that were transiently transfected with optiFIX and non-transfected cells (WT), respectively. **e** Immunodetection of extracellular FIX protein with anti-FIX Ab (1:5,000). Culture supernatant of non-transfected cells was used as negative control (Neg). T1, T2: Two different transient transfections with the optiFIX plasmid. M: 1 kb Marker (Thermo Fisher Scientific).

Next, we compared the codon usage bias in *P. patens* to biases of other pharmaceutical production platforms. The codon usage frequencies of under- and over-represented codons for Cys, Glu, Phe, His, Lys, Asn, Gln, and Tyr in *P. patens* were compared with *Spodoptera frugiperda* (insect cells), *Homo sapiens* (HEK cells), *Cricetulus griseus* (CHO cells), *Nicotiana tabacum* (tobacco) and *Oryza sativa* (rice). For instance, the over-represented codon for Cys, TGC, in *P. patens* (9.8 in thousands) is also an over-represented codon in *S. frugiperda*, *H. sapiens*, *C. griseus*, and *O. sativa*. On the other hand, this codon is under-represented in *N. tabacum* and TGT is preferred over TGC for Cys. The synonymous codon usage bias in *P. patens* is similar to codon usage biases in *S. frugiperda*, *H. sapiens*, *C. griseus* (CHO cells), and *O. sativa* (rice), but different from *N. tabacum* (tobacco) (Table 1).

**Table 1:** The codon usage frequencies of under- and over-represented codons for eight amino acids in different protein expression systems. Numbers represent the frequency of codon per thousand. Data are from http://www.kazusa.or.jp/codon/.

*Codon frequency in thousands*
|  |  | *Physcomitrella patens* | *Spodoptera frugiperda* | *Homo sapiens* (HEK cells) | *Cricetulus griseus* (CHO cells) | *Oryza sativa* | *Nicotiana tabacum* |  |
| --- | --- | --- | --- | --- | --- | --- | --- | --- |
| *under-represented* | TGT | 7.0 | 8.5 | 10.6 | 9.1 | 6.2 | 9.8 | Cys |
| *over-represented* | TGC | 9.8 | 13.2 | 12.6 | 10.3 | 12.4 | 7.2 |  |
| *under-represented* | GAA | 24.1 | 27.6 | 29.0 | 28.4 | 21.6 | 36.0 | Glu |
| *over-represented* | GAG | 37.4 | 33.1 | 39.6 | 41.1 | 38.6 | 29.4 |  |
| *under-represented* | TTT | 17.2 | 10.1 | 17.6 | 19.6 | 13.1 | 25.1 | Phe |
| *over-represented* | TTC | 23.6 | 27.5 | 20.3 | 22.0 | 22.4 | 18.0 |  |
| *under-represented* | CAT | 11.1 | 8.7 | 10.9 | 10.2 | 11.3 | 13.4 | His |
| *over-represented* | CAC | 11.6 | 15.6 | 15.1 | 12.9 | 13.8 | 8.7 |  |
| *under-represented* | AAA | 18.9 | 26.8 | 24.4 | 24.6 | 16.0 | 32.6 | Lys |
| *over-represented* | AAG | 34.7 | 49.2 | 31.9 | 38.4 | 32.3 | 33.5 |  |
| *under-represented* | AAT | 17.8 | 13.4 | 17.0 | 17.4 | 15.1 | 28.0 | Asn |
| *over-represented* | AAC | 20.4 | 28.8 | 19.1 | 21.2 | 18.5 | 17.9 |  |
| *under-represented* | CAA | 16.7 | 16.1 | 12.3 | 10.3 | 13.5 | 20.7 | Gln |
| *over-represented* | CAG | 22.0 | 21.6 | 34.2 | 33.4 | 20.8 | 15.0 |  |
| *under-represented* | TAT | 10.5 | 10.0 | 12.2 | 13.1 | 10.0 | 17.8 | Tyr |
| *over-represented* | TAC | 17.2 | 24.4 | 15.3 | 16.4 | 15.1 | 13.5 |  |

### Comparison of protein levels of FIX and optiFIX *in vivo*

In order to analyze whether the protein amounts of FIX and optiFIX show any difference in live moss cells, we cloned two fusion constructs, FIX-Citrine and optiFIX-Citrine. Subsequently, we transiently transfected moss protoplasts with these constructs, and performed confocal laser scanning microscopy (CLSM) on day 3 after transfection. A visual analysis of the images revealed that the FIX-Citrine signal shows a diffuse localization pattern, which might be due to intracellular mislocalization of aberrant protein isoforms (Fig. 6a). In contrast, optiFIX-Citrine exhibited a complex network-like organization, indicating a proper localization of the protein in the ER (Fig. 6b). Furthermore, the optiFIX-Citrine signal intensity was higher than that of FIX-Citrine. Comparison of mean voxel intensities calculated for FIX-Citrine and for optiFIX-Citrine (3.88 and 41.73, respectively) shows a nearly eleven-fold higher fluorescence intensity after the codon optimization, which had prevented undesired heterosplicing (Fig. 6c).

**Fig. 6:**
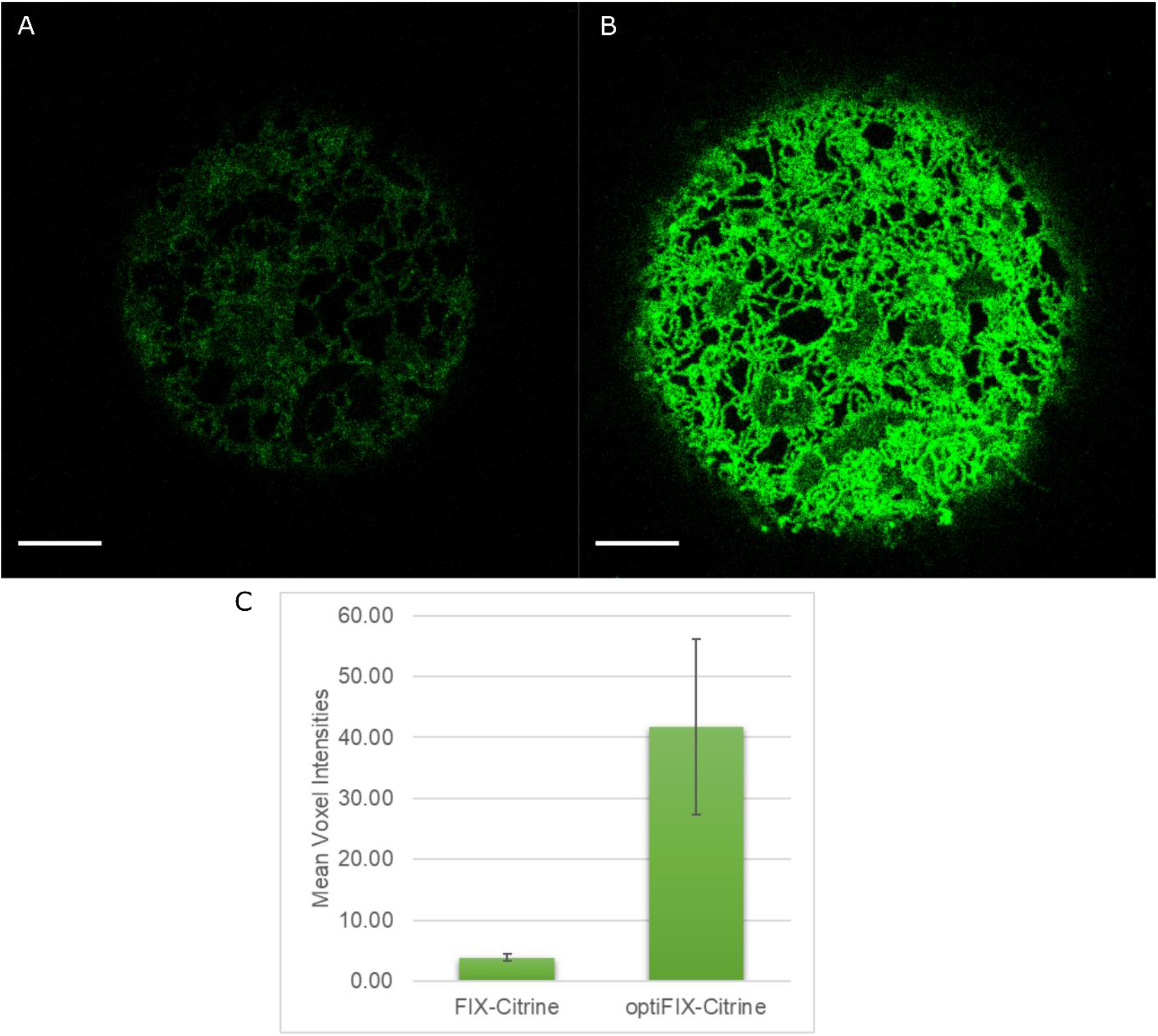
Comparison of FIX and optiFIX protein levels using confocal microscopy and quantitative image analysis. **a** Transient overexpression of FIX-Citrine. The slice with the brightest mean intensity was extracted from a Z-stack and its voxel intensity range was adjusted to (0-36) for illustration. Scale bar is 5 µm. **b** Transient overexpression of optiFIX-Citrine. The slice with the brightest mean intensity was extracted from a Z-stack and its voxel intensity range was adjusted to (0-36) for illustration. Scale bar is 5 µm. **c** Comparison of the mean signal intensities of the Z-stacks for FIX-Citrine and optiFIX-Citrine (n=3).

### Automatic codon-optimization of transgenic coding sequences

In order to automatize the process of *P. patens*-specific codon optimization, we compiled our findings into a JavaScript-based web application: physCO, and made it available at www.plant-biotech.uni-freiburg.de. physCO takes as input coding sequences in FASTA format, inspects them codon-wise, substitutes codons where possible according to our findings on codon usage (Table 2) and finally outputs them in FASTA format. In addition, physCO checks the input for consistent lengths, valid start and stop codons and internal, premature stop codons and raises warnings in case it detects irregularities.

**Table 2:** Under-represented codons were changed into the over-represented ones based on the codon usage tables ^49,50^ to optimize FIX sequence using aspFIX as a template.

| **Amino acid** | **Under-represented codons in aspFIX** | **For optiFIX, changed into** |
| --- | --- | --- |
| **Cys** | TGT | TGC |
| **Glu** | GAA | GAG |
| **Phe** | TTT | TTC |
| **His** | CAT | CAC |
| **Lys** | AAA | AAG |
| **Asn** | AAT | AAC |
| **Gln** | CAA | CAG |
| **Tyr** | TAT | TAC |

### Splicing of FH mRNA like FIX occurs in transgenic plants

In order to test whether heterosplicing is a common phenomenon in *P. patens* upon transgene expression, we analyzed already existing recombinant human complement factor H (FH)-producing moss lines^40^. Our RT-PCR analysis revealed that expression of the human FH CDS in moss generated two transcript variants, the full-length FH transcript of 3,639 bp and a 1,215 bp FH variant, which could lead to a truncated FH isoform with a molecular mass of 45.6 kDa (Supplementary Fig. 3). Therefore, we generated with the help of physCO an optimized version of hFH, and named it optiFH. These changes affected 386 of 1213 codons, and increased the GC% content from 33.9% to 49.8% (Supplementary Fig. 4).

To investigate the effects of these alterations in live moss cells, we cloned two fusion constructs, FH-Citrine and optiFH-Citrine. Subsequently, we transiently transfected moss protoplasts with these constructs, and performed CLSM on day 3 after transfection. A visual analysis of the images acquired for FH and optiFH revealed that while an FH-Citrine signal was not detectable, optiFH-Citrine signal intensity was high and filled the secretory pathway (Supplementary Figure 5). Analysis on FH showed that heterosplicing is not an effect specific for FIX, but a general phenomenon in moss. Therefore, transgene sequences have to be modified using physCO prior to transfection for proper transcription and translation.

## DISCUSSION

Heterologous gene expression is of fundamental importance for basic biology and for industrial production. In order to optimize transgene transcription and translation as well as transcript stability, many factors have to be taken into account. Engineering of upstream and downstream regulatory sequences, replacing original signal peptides with host-suitable ones, and optimizing codons according to host codon-usage pattern can be used to improve protein production.

Splicing/alternative splicing is another important factor that alters expression rates by the generation of various transcripts, which create different protein isoforms. Generation of these protein isoforms can affect fundamental characteristics of the protein such as function, activity, intracellular localization, interaction with other molecules, and stability. Moreover, splicing can create a transcript with an early stop codon, which may be degraded by NMD. Thus, it can be hypothesized that modulation of splicing, a neglected point in transgene expression, can improve protein production. For instance, the expression of GFP from the jellyfish *Aequorea victoria* in *A. thaliana* failed due to aberrant mRNA processing^30^. To avoid AS, generally cDNAs derived from already spliced mRNAs are introduced into the genome of the production host.

We introduced the human FIX cDNA under the control of the *PpActin5* promoter and CaMV 35S terminator into the *P. patens* genome. RNA was isolated from protonemata of transgenic lines and RT-PCR analysis revealed unexpectedly that a fraction of human FIX mRNA was spliced into four variants in addition to the unspliced full-length transcript, resulting in five mRNAs a different lengths derived from one cDNA. According to RT-PCR analyses, this phenomenon occurred not only in juvenile protonema, but also in adult moss plants. We sequenced these mRNAs and surprisingly found that reading frames had not changed in the four smaller variants in comparison to the mature full-length transcript. This may have happened by a pure coincidence. Alternatively, we may not have been able to detect such putative minor transcripts because they were either too low abundant for RT-PCR, or they have been eliminated by NMD.

The outcome of this heterosplicing was detectable on four levels, by RT-PCR from mRNAs, immunoblot, mass spectrometry of the protein isoforms, and *in vivo* by confocal imaging showing deviant localization of Citrine reporter fusions. These findings were also reflected in RT-PCR analysis from transiently transfected moss cells. Our observations revealed that heterosplicing of the human FIX mRNA in moss is consistent across different tissues and not dependent on developmental stages. Furthermore, the consistency of heterosplicing in episomally (transiently) FIX-expressing cells indicates that heterosplicing of the FIX mRNA in transgenic lines was neither due to illegitimate or partial integration of the construct into the moss genome nor a locus-specific effect. Moreover, results of PCRs from genomic DNA of transgenic lines supported this conclusion. Unlike a previously reported case, in which expression of full-length GFP transcript and fluorescence were not detectable^30^, we report here for the first time heterosplicing of a human cDNA in a plant. It caused the generation of various FIX mRNA variants and protein isoforms in addition to the complete transcript and protein. Therefore, a possible occurrence of heterosplicing should be analyzed on RNA level, even if a full-length protein can be detected in heterologous cells.

All splice junctions in the FIX CDS were determined by sequencing. In addition, we identified consensus motifs of donor and acceptor splice sites in the latest *P. patens* genome version^45^ using all 87,533 annotated transcripts that originate from all 32,926 protein-coding genes, and compared them with our experimentally verified motifs. This revealed that *P. patens*, like most eukaryotes, follows the GT-AG splicing rule. The same was true for the cryptic additional introns recognized in the FIX mRNA; obviously, the *P. patens* spliceosome interprets parts of the FIX CDS as intronic sequences. Presumably, moss cells recognize motifs in the human FIX CDS as *cis* exonic and intronic splicing enhancers. Little is known about exonic and intronic regulatory sequences in plants^51,52^. Moreover, plant introns are different from animal introns in terms of UA- or U-richness, which is crucial for splicing efficiency^53^. Due to these characteristic differences, it is likely that the GC content can affect the characteristics of splice site recognition^15^. The GC content is correlated with many features including gene density^54^, intron length^55^, meiotic recombination^56^, and gene expression^57^. It varies among species, and even along chromosomes^58^. It was shown before that the average GC content in *P. patens* coding sequences is 50%^46^. The GC content of the human FIX CDS recognized as cryptic additional introns ranged from 38% to 40%. This remained the same after exclusively mutating the splice sites’ environment in aspFIX; even though mutations in the splicing motifs decreased the number of transcripts from five to two, the GC content of the cryptic alternative intron was still only 40%. The shorter transcript was not in frame, but still detectable by RT-PCR. Therefore, we created the fully codon-optimized plasmid optiFIX to check the effect of overall codon optimization on pre-mRNA splicing. A closer comparison of the aspFIX and optiFIX constructs revealed that there was no change in the donor and acceptor sites of the newly emerged cryptic intron. There was, however, an increase of the GC content within the cryptic intron sequence. The GC contents of the sequences previously defined as introns in the FIX and aspFIX constructs increased to 46% - 50% in optiFIX. Hence, we conclude that the increase of GC content introduced by the overall codon optimization together with mutating splice siteś environment prevented the heterosplicing, the mis-interpretation of any sequence as intron by the moss spliceosome. As a result, a nearly eleven-fold increase in protein amounts was achieved in transiently transfected moss protoplasts *in vivo.* As heterosplicing did not cause detectable frameshift mutations, fluorescence signals obtained from the FIX-Citrine construct were the sum of all FIX protein isoforms, full-length and fragmentary, fused to Citrine. Based on this inference and proteins, which were harvested from transient transfection with the FIX construct, and detected on the immunoblot (Fig. 4), we conclude that the prevention of heterosplicing from the optiFIX construct has improved full-length protein amounts *in vivo* even more than eleven-fold.

Two mechanisms, namely boosted translation rates and the prevention of mRNA degradation via NMD, may contribute to this result. In addition, the protein from the optiFIX construct was solely detectable in the ER, the compartment of choice for correct post-translational modifications and subsequent secretion of the protein. A similar phenomenon was observed in the expression of another transgene in *P. patens.* One example of heterosplicing and its prevention by our methodology is the expression of human complement factor H (FH). FH is an important regulator of the alternative pathway in the human complement system. Currently, a recombinant FH is not available on the market, although it has potential as a biopharmaceutical in the treatment of severe human diseases like atypical hemolytic uremic syndrome (aHUS), age-related macular degeneration (AMD) or C3 glomerulopathies (C3G). The recombinant production of FH, devoid of plant-specific N-linked sugar residues, in *P. patens* resulted in a range of promising biological activities^40^. Moreover, moss-derived FH is currently being examined for use in Covid-19 treatment (www.elevabiologics.com). RT-PCR analysis in FH-producing stable moss lines revealed that the expression of human FH CDS generated 2 transcript variants, the full-length FH transcript of 3,639 bp and a 1,215 bp FH variant that could lead to the production of an FH isoform with a calculated molecular mass of 45.6 kDa (Supplementary Fig. 3). We optimized the FH CDS using our bioinformatics tool physCO, described here, to analyze whether the protein amounts of FH and optiFH show any difference in live moss cells. A visual analysis of the images acquired for FH and optiFH from transiently transfected moss cells revealed that while FH-Citrine signal intensity is not detectable, optiFH-Citrine signal intensity is high (Supplementary Fig. 5). Thus, an optimization of the currently used CDS of human FH in plants is advantageous for plant-based production.

These findings suggest that the splicing of heterologous mRNAs should be taken into account on a routine basis. In addition, our analysis of synonymous codon usage bias suggests that our methodology to prevent heterosplicing can be directly implemented in organisms, which follow similar codon usage patterns as *P. patens*, such as *O. sativa*, insect or mammalian cells. Our methodology might still be vital in organisms that have different codon usage patterns, such as the species of the genus *Nicotiana*, e.g. *N. tabacum* and *N. benthamiana.* Mutating donor and acceptor sites and their neighboring nucleotides together with replacing codons ending with A or T to G or C may prevent the generation of heterosplice variants.

Why should splice site mutation and codon optimization be employed at the same time rather than codon optimization alone? Although differences between GC-rich and GC-poor genes were not reported in moss, analyses of various plant species can explain why codon optimization might not be sufficient. The GC content of GC-poor CDS in *A. thaliana*, soybean, pea, tobacco, tomato, and potato ranges from 40-46% while GC content of introns is 26-31%^59^. A similar trend is visible in GC-rich genes: 46-49% in CDS and 29-31% in introns. On the other hand, in maize, the GC content is higher: 56% in GC-poor CDS and 67% in GC-rich CDS. The GC content of introns in maize is also elevated: 40% in GC-poor genes and 48% in GC-rich genes^59^. This shows that the intron/exon definition is not solely dependent on the GC content. Hence, increasing the GC content of cryptic introns to a certain level without mutating splice site motifs might not be sufficient to prevent aberrant mRNA processing.

In general, the analysis of spliced transcripts is beneficial for at least five reasons. (i) It facilitates the reliable quantification of gene expression by designing primers that differentiate between full-length transcripts and shorter variants. (ii) It enables the precise detection and quantification of heterologous proteins by antibodies that differentiate functional isoforms from non-functional ones. (iii) It helps to better purify proteins either by specific antibodies or by size exclusion chromatography. (iv) Optimized cDNAs may result in better detectable reporter proteins *in vivo*. (vi) Aberrant protein isoforms due to heterosplicing may show mislocalization in cells of the production host and thus may contribute to poorly modified glycoproteins. Prevention of heterosplice variants eventually will improve recombinant protein production and decrease downstream processing costs significantly. Hence, plant-based systems can become an alternative to traditional production platforms. Moreover, it may have implications for basic biology as well, because here the use of heterologous reporter constructs is of vital importance. For these purposes, we developed physCO, an automated *P. patens*-specific codon-optimization tool, which allows the optimization of transgene expression by the prevention of undesired mRNA splicing. Moreover, our analysis of codon usage patterns indicates that this tool can also be used for insect (*S. frugiperda),* HEK (*H. sapiens)*, and CHO (*C. griseus)* cells as well as rice (*O. sativa)*. Besides that, we cannot exclude the possibility, that the phenomenon of mRNA splicing here described as heterosplicing is a novel gene regulatory mechanism occurring in eukaryotes in general.

## MATERIALS AND METHODS

### Design of the expression vectors pFIX, aspFIX, and optiFIX

The coding DNA sequence for human Factor IX without the native signal peptide was synthesized as N-terminal translational fusion to the signal-peptide coding sequence of PpAP1 (Pp3c5_19520V3.1)^42^ by GeneArt (Thermo Fisher Scientific, Waltham, MA, USA). It was cloned into the pAct5-MFHR1 plasmid^41^ to generate the final FIX expression vector, pFIX. Its expression is driven by the promoter of the *PpActin5* gene^60^ and the CaMV 35S terminator.

Nucleotides involved in splicing were replaced without a change of the FIX aa sequence in the aspFIX plasmid which was synthesized by GeneArt. For codon optimization, two sources, codon usage bias calculated by Hiss *et al.*^49^ and Kazusa codon usage database^50^ (http://www.kazusa.or.jp/codon/cgi-bin/showcodon.cgi?species=145481), were used. Unlike a recent report^49^, we decided to exchange the eight under-represented codons for Cys, Glu, Phe, His, Lys, Asn, Gln and Tyr. Under-represented codons were replaced with the over-represented alternative codons (Table 2) via physCO. The optiFIX CDS was synthesized by GeneArt. See Fig. 7 for a multiple sequence alignment of FIX, aspFIX, and optiFIX.

**Fig. 7:**
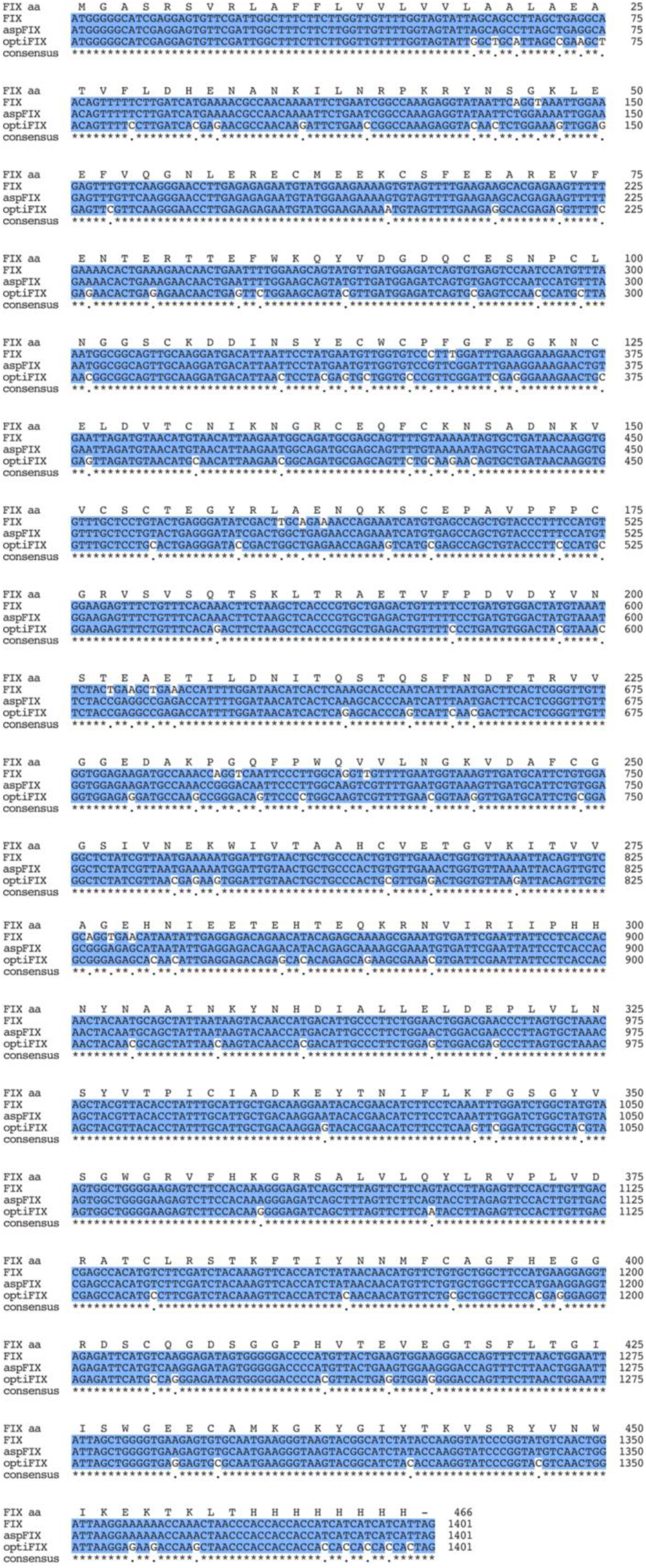
Multiple sequence alignment of FIX, aspFIX and optiFIX CDS. The translated amino acid sequence of FIX (FIX aa) was added for better visualization. Alignment was performed and formatted for publication with MUSCLE via the “MSA” package for R^49,50^.

### Design of the expression vectors pFH and optiFH

The previously generated Factor H expression plasmid pFH^40^ contains the same promoter and terminator as pFIX. It was used in transient expression assays. The optiFH CDS was deduced via physCO and synthesized by GeneArt. It was cloned into the same vector as FH. See Supplementary Fig. 4 for a sequence alignment of FH and optiFFH.

### Transfection and screening of transgenic plants

The *Physcomitrella patens* (Hedw.) B.S. Δ*xt/ft* moss line, a double knockout line in which the *α1,3 fucosyltransferase* and the *β1,2 xylosyltransferase* genes had been abolished^61^ (IMSC accession number 40828), was cultured in liquid Knop medium (pH 4.5) one week prior to transfection as described previously^62^ to obtain cells (protoplasts). Protoplast transfection was performed as described^63^ using 50 µg of linearized pFIX DNA per transfection. In addition to the generation of transgenic lines, mossFIX was produced transiently by an upscaled transfection protocol based on the proportion of 0.7-1 µg DNA of pFIX for 1×10^5^ cells. Stable transfectants were selected on solidified Knop medium containing 25 µg/mL hygromycin as described^62^. Cells transiently expressing mossFIX were grown in special regeneration medium (7.5 mM MgCl_2_×6H_2_O, 2.6 mM MES, 200 mM D(−)-Mannitol, 62.6 mM CaCl_2_×2H_2_O, 68.4 mM NaCl, 2.8 mM D(+)-Glucose monohydrate, 5 mM KCl; pH: 5.6; osmolarity: ~550–570 mOsm) and harvested for RNA and protein analyses two weeks after transfection. To analyze whether splicing of FIX was prevented on RNA level, WT cells were also transfected with the aspFIX and optiFIX constructs, respectively, and harvested for RNA isolation two weeks after transfection.

### Plant cell culture

After selection on solidified Knop medium, transgenic lines were cultured under standard conditions in liquid Knop medium^64^. They were subcultured periodically by disrupting the tissue with the use of an Ultra-Turrax (IKA, Staufen, Germany) at a rotational speed of 16,000 – 18,000/min for 1 min. Subsequently, they were transferred to fresh medium.

### Molecular characterization of transgenic lines

Following the hygromycin selection process, stable lines were characterized by the presence of the FIX transcript. To validate the presence of the transgene in the moss genome, 80-100 mg (fresh weight) moss material from protonema cultures were disrupted with the use of a stainless steel ball in TissueLyser II (Qiagen, Hilden, Germany) for 1 min with an impulse frequency set to 30. Genomic DNA was extracted using GeneJET Genomic DNA Purification Kit (Thermo Fisher Scientific, Waltham, MA, USA) according to the manufacturer’s instructions. Transgene presence and completeness were verified with a standard PCR with the primers FIXfwdB (5’-GGAGTGTTCGATTGGCTTTCT-3’) and FIXrevB (5’-TGGTGGTGGTGGGTTAGTTT-3’) that amplifies almost the full-length FIX CDS. For transgene expression analysis via reverse transcription (RT)-PCR, 50-100 mg (fresh weight) moss material from protonema cultures were disrupted with the use of a stainless steel ball in TissueLyser II for 2 min with an impulse frequency set to 30. The material was resuspended in 1 mL TRIzol^®^ Reagent (Thermo Fisher Scientific), and RNA was isolated according to the manufacturer’s instructions. For analysis of gametophores, 5-6 gametophores were collected and RNA isolation was performed as described above. Six micrograms of DNase-treated RNA were used for first strand synthesis with Superscript Reverse Transcriptase III (Thermo Fisher Scientific). The quality of cDNA was controlled with a standard PCR with the primers c45for (5’-GGTTGGTCATGGGTTGCG-3’) and c45rev (5’-GAGGTCAACTGTCTCGCC-3’) corresponding to the gene coding for the ribosomal protein L21. Transgene expression was verified with the primers FIXfwdB and FIXrevB that amplifies almost the full-length FIX CDS. PCR reactions were performed with Phusion high-fidelity DNA polymerase (Thermo Fisher Scientific). The PCR products were examined by standard agarose gel electrophoresis with visualization of DNA by ethidium bromide fluorescence.

### Molecular characterization of transiently transfected cells

Following the transfection, cells were grown in regeneration medium for two weeks^65^. Transiently transfected cells (nearly 3.6 million cells for FIX and aspFIX; 7.2 million cells for optiFIX) were used for transgene expression analysis via RT-PCR. RNA was isolated using the TRIzol^®^ Reagent and RNA was purified through spin columns of Direct-zol™ RNA MicroPrep kit (Zymo Research, Irvine, California, USA) according to the manufacturer’s instructions. After DNase-treatment and first-strand synthesis (for the cDNA synthesis: 0.4 µg RNA were used for FIX and aspFIX; 2 µg RNA were used for optiFIX), the quality of cDNA was controlled with primers c45for and c45rev. Transgene expression was verified again with primers FIXfwdB and FIXrevB.

### Analysis of FIX cDNA products

Amplified products were excised from the gel and purified using QIAEX II Gel Extraction Kit (Qiagen). Sequencing was done by Eurofins Genomics (GATC Sanger sequencing service). In the sequencing reactions, in addition to FIXfwdB and FIXrevB, FIX6deepseq1F (ATGGGGGCATCGAGGAGTGTTCGAT), FIX6deepseq1R (TTCCCCAGCCACTTACATAGCCAGA), and FIXseq6 (CACCAACAACCCGAGTGAAG) were used. The remaining purified products were separately cloned into the pJET vector using the CloneJET PCR Cloning Kit (Thermo Fisher Scientific) according to the manufacturer’s instructions. Clones with aberrant insert sizes were selected for plasmid isolation with the GeneJet Plasmid Miniprep kit (Thermo Fisher Scientific). Sequencing was done as described above.

### Splicing motif search

The most recent release of the *Physcomitrella* patens v3.3 genome annotation^45^, which is available from https://phytozome.jgi.doe.gov/, was used. The splice sites of all 87,533 annotated transcripts from all 32,926 protein-coding genes were used as anchors to extract surrounding genomic sequences with a fixed window size of ± 9 bp. Consensus sequence logos of all extracted donor and acceptor sites were created using R and the package ggseqlogo^66,67^.

### Codon usage bias among different species

Synonymous codon usage biases in *P. patens, S. frugiperda, H. sapiens, C. griseus, O. sativa,* and *N. tabacum* were obtained from the Codon Usage Database (https://www.kazusa.or.jp/codon/) and biased codons in moss were compared with the codon usage pattern of other species.

### Confocal microscopy imaging and image analysis

All images were taken with a Leica TCS SP8 microscope (Leica Microsystems, Wetzlar, Germany) using HCX PL APO 63x/1.40 oil objective with a zoom factor of 4.5. For the excitation of Citrine, an WL laser was applied at 2% with an excitation wavelength of 514 nm. The voxel sizes were 0.080 μm on the X-Y dimensions and 0.300 μm on the Z dimension. The pinhole was adjusted to 1 Airy Unit (95.5 μm).

The image processing consists of the following steps: i) denoising, ii) edge enhancement, iii) Richardson-Lucy restoration, iv) local intensity equalization, v) segmentation. In the first step, a median filter with a window of (3,3,3) was applied to the images in order to remove the salt-and-pepper noise. In the second step, the denoised images were subjected to an unsharp-mask operation^68,69^, which involves blurring the image with a Gaussian filter, subtracting the blurred image from the input image, and adding the resulting difference back to the input image, a process that sharpens the edges in the images. The edge-enhanced images were then subjected to the Richardson-Lucy restoration algorithm, as implemented in the scikit-image package^70^ for three iterations, assuming an averaging filter in a window of (3,5,5) as point spread function (here the aim was to smooth the image without losing the thin structures, rather than a true deblurring). The code for the Richardson-Lucy algorithm was adapted from scikit-image package^70^ using Python programming language. Subsequently, a spatial intensity equalization step was implemented on the images using a home-built algorithm. In short, each image was divided into boxes with dimensions of (7,7,7) voxels. The voxel values within these boxes were rescaled by multiplying the values in each box with weights that are calculated based on the skewness value corresponding to the same box. This step corrected for the intensity gradients in the image in order to minimize the loss of some low-intensity foreground voxels. In the last step, the images were segmented by applying an adaptive thresholding method, where the thresholds were determined locally by applying Otsu’s method to local windows of (7,19,19) voxels throughout the image volume. The code for the adaptive Otsu thresholding algorithm was developed based on the Otsu threshold function of the scikit-image package^70^ and the Numba package^71^. To suppress over-segmentation of noisy background areas, a low global threshold was also applied by first calculating Otsu’s threshold for whole image and multiplying it with 0.66. The adaptive thresholding operation yielded the binary masks that specified the foreground voxels, which in turn were used to calculate the mean voxel intensity for each original image. Three replicates were used for analysis of each of the FIX-Citrine and optiFIX-Citrine constructs. The same is true for FH-Citrine and optiFH-Citrine but the quantification was not possible due to undetectable/very low signals in FH-Citrine images. All six FIX images were processed and quantified in a single run of a Python script by using the exact same parameters to avoid any possible bias. Other packages used to develop the code are: NumPy^70,72^, Pandas^73^ and SciPy^74^. The code used for the image processing and quantification can be obtained from www.plant-biotech.uni-freiburg.de. The procedure is shown in Supplementary Fig. 6.

### Recombinant protein extraction and detection

Proteins from the culture supernatant were precipitated as described^75^. After air drying, the precipitate was resuspended in 50 mM HEPES, 2% SDS (pH 7.5) and disulfide bonds within proteins were reduced with 25 mM dithiothreitol (DTT), followed by alkylation of the thiol groups with 60 mM iodoacetamide (IAA). For Western blot analyses, 7.5% SDS polyacrylamide gel electrophoresis (SDS-PAGE Ready Gel^®^ Tris-HCl Precast Gels, BioRad, CA, USA) was run at 100 V for 1h 30min and blotted to polyvinylidene fluoride (PVDF) membranes (Amersham Hybond P 0.45 µm pore size PVDF blotting membrane, GE Healthcare) in a Trans-Blot^®^ SD Semi-Dry Transfer Cell (BioRad) for 2 h with 1.8 mA/cm^2^ membrane. The membrane was blocked for 1 h at room temperature in TBS containing 4% Amersham ECL Prime Blocking Reagent (GE Healthcare) and 0.1% Tween 20. It was then incubated with polyclonal anti-factor IX antibody (F0652, SIGMA) (1:3000) overnight at 4°C. Afterwards, the membrane was washed three times with TBS containing 0.1% Tween 20 and incubated with horseradish peroxidase-linked anti-rabbit IgG (NA934, GE Healthcare) at a dilution of 1:10,000 for 1 h. The blot was washed again and proteins were detected using the Amersham ECL Prime Western Blotting Detection Reagent (GE Healthcare) according to the manufacturer’s instructions.

### Sample preparation and MS analysis

The reduced and alkylated protein extracts were subjected to SDS-PAGE. Appropriate gel bands showing FIX fractions were excised and processed for in-gel digestion: Following the destaining of the gel bands with 30% acetonitrile (ACN), they were treated with 100% ACN and dried completely in a vacuum concentrator. Tryptic in-gel digestion of gel slices was performed overnight at 37°C in 50 mM ammonium bicarbonate using 0.1 µg trypsin (Promega) per gel band. Afterwards, peptides were extracted from the gel with 5% formic acid. Analyses were performed using the UltiMate 3000 RSLCnano system (Dionex LC Packings/Thermo Fisher Scientific) coupled online to a QExactive Plus instrument (Thermo Fisher Scientific). The UHPLC systems was equipped with a C18-precolumn (Ø: 0.3 mm, 5 mm; PepMap, Thermo Fisher Scientific) and an Acclaim^®^ PepMap analytical column (ID: 75 m, 500 mm, 2 m, 100 Å, Dionex LC Packings/Thermo Fisher Scientific).

MS analyses were performed using a binary solvent system consisting of 0.1% formic acid (FA, solvent A, “A”) and 0.1% FA/86% ACN (solvent B, “B”). Samples were washed and concentrated on a C18 pre-column with 0.1% TFA for 5 min before switching the column in line with the analytical column. Peptide separation was performed applying a 45-min gradient at a flow rate of 250 nl/min. Peptide samples were eluted with a gradient of 4–40% B in 30 min and 40–95% B in 5 min. After each gradient, the analytical column was washed with 95% B for 5 min and re-equilibrated for 15 min with 4% B. The MS instrument was externally calibrated using standard compounds and equipped with a nanoelectrospray ion source and distal coated SilicaTips (FS360–20-10-D, New Objective, Woburn, MA, USA), and MS/MS analyses were performed on multiply charged peptide ions. The measurement was performed using UltiMate 3,000 RSLCnano system (Dionex LC Packings/Thermo Fisher Scientific, Dreieich, Germany) coupled online to a QExactive Plus mass spectrometer (Thermo Fisher Scientific, Bremen, Germany) and mass spectrometer was operated in the data-dependent mode to automatically switch between MS (max. of 1×10 ions) and MS/MS. Each MS scan was followed by a maximum of 12 MS/MS scans using HCD with a normalized collision energy of 35%. The mass range for MS was m/z = 375–1,700 and resolution was set to 70,000. MS parameters were as follows: spray voltage 1.5 kV; ion transfer tube temperature 200°C. Raw data were analyzed using Mascot Distiller V2.5.1.0 (Matrix Science, USA) and the peak lists were searched with Mascot V2.6.0 against an in-house database containing all *P. patens* V1.6 protein models^22^ as well as human FIX (complete and alternative ones based on splicing). The peptide mass tolerance was set to 8 ppm and the fragment mass tolerance was set to 0.02 Da. carbamidomethylation (C, +57.021464 Da) was specified as fixed modification. Variable modifications were Gln->pyro-Glu (−17.026549 Da), oxidation/hydroxylation (M, P, +15.994915 Da) and deamidation (N, +0.984016 Da). A total of two missed cleavages was allowed as well as semitryptic peptides. Search results were loaded into Scaffold4 software (Version 4.11.0, Proteome Software Inc., Portland, OR) and an additional database search using X!Tandem^76^ implemented in the software on the loaded spectra was performed.

## ACKNOWLEDGEMENTS

We thank Bettina Warscheid for the possibility to use the QExactive Plus mass spectrometer and Anne Katrin Prowse for language editing. This work was supported by the Excellence Initiative of the German Federal and State Governments (GSC-4 to O.T., EXC-294 and EXC-2193/1 – 390951807 to R.R.); and a grant from the German Federal Ministry of Education and Research (BMBF 0313852C to R.R.).

## AUTHOR CONTRIBUTIONS

O.T., E.L.D. and R.R. designed research; O.T., S.W.L.M., B.Ö., and N.v.G. performed research and together with S.N.W.H analyzed data; O.T., E.L.D., and R.R. wrote the paper.

## DATA AVAILABILITY

All data generated or analyzed during this study are included in this article and its supplementary information.

## CODE AVAILABILITY

JavaScript-based web application, physCO, as well as the code used for the image processing and quantification are available at www.plant-biotech.uni-freiburg.de.

## Supplementary Data

**Supplementary Fig. 1:**
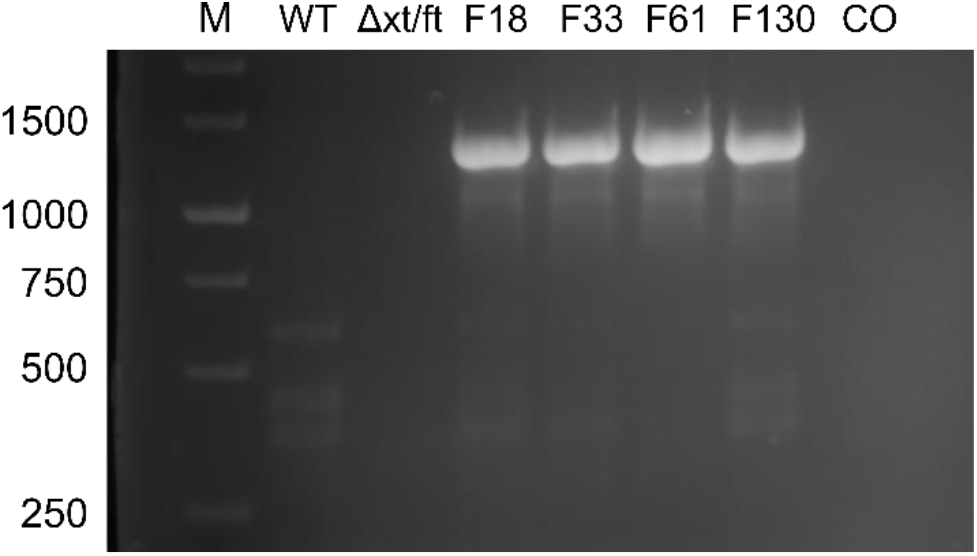
The presence of complete FIX CDS in transgenic lines. Genomic DNA from protonema cultures of 4 FIX-transgenic lines (F), the parental line Δ*xt/ft*, and WT was prepared and used for PCR using the primers FIXfwdB and FIXrevB. M: 1 kb Marker (Thermo Fisher Scientific), CO: Water control.

**Supplementary Fig. 2:**
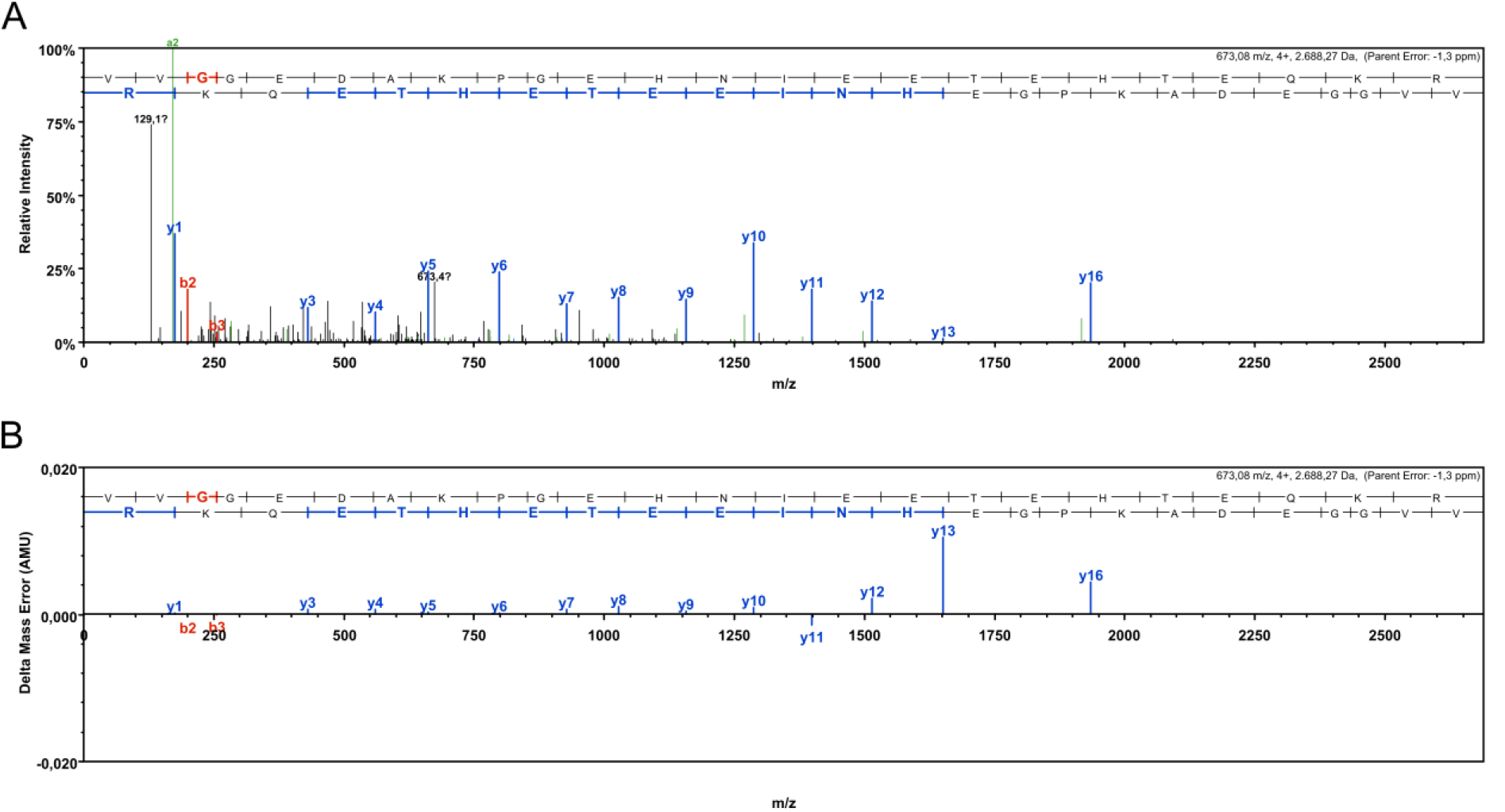
Identification of the peptide VVGGEDAKPGEHNIEETEHTEQKR by MS. **a** HCD fragment ion spectrum of the identified peptide. **b** Fragment mass error distribution of the b- and y-ion series.

**Supplementary Fig. 3:**
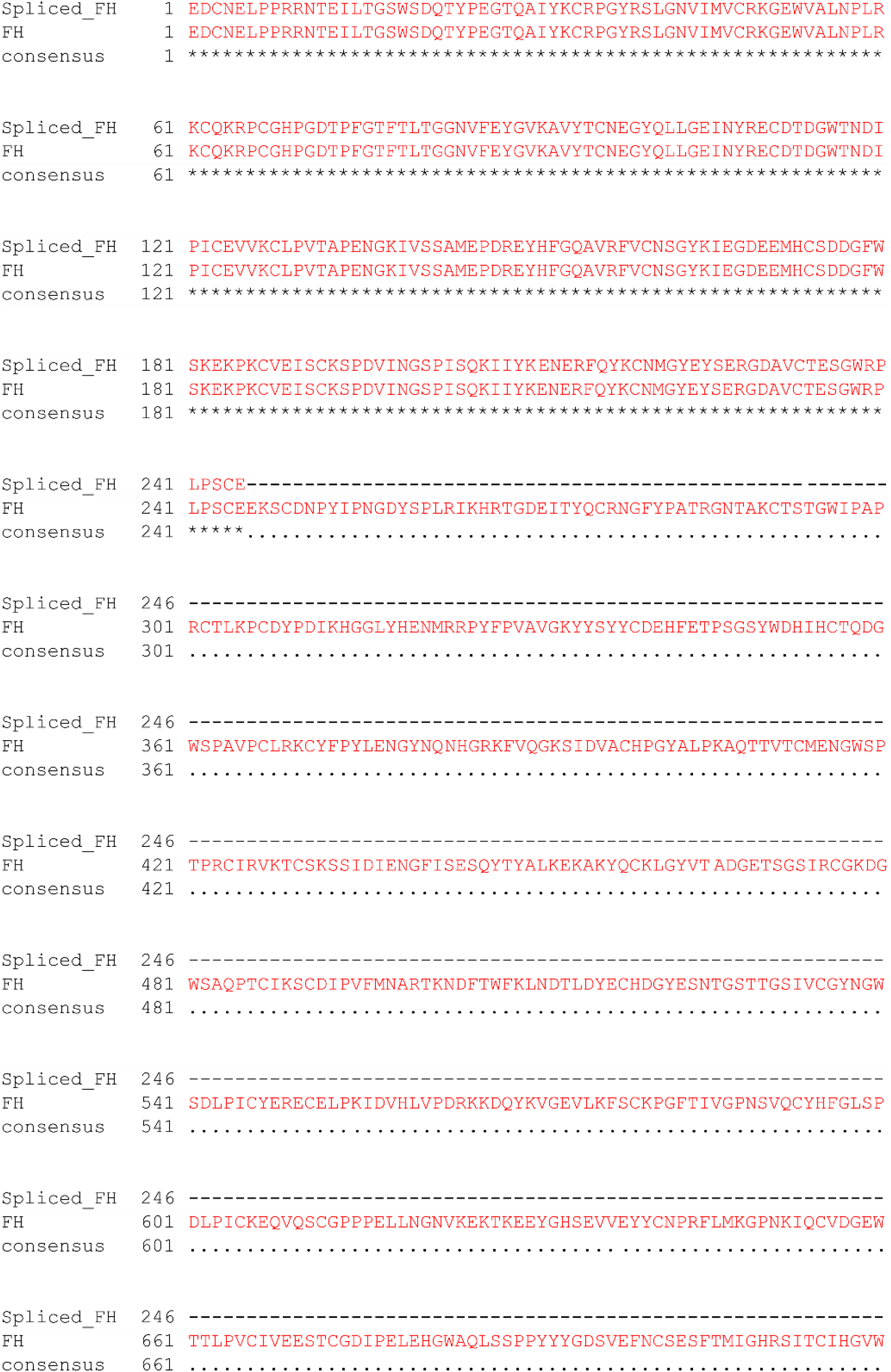
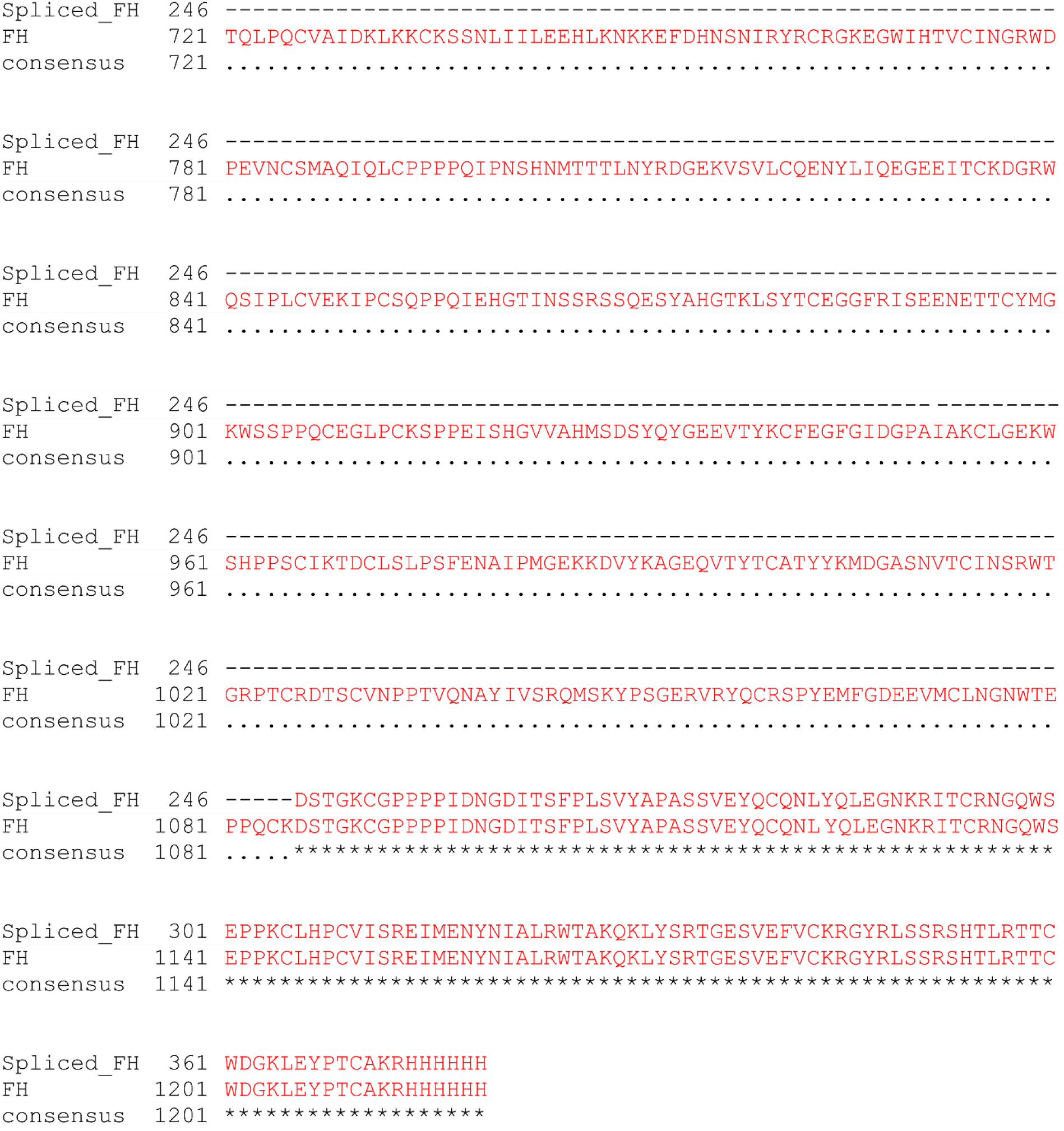
Pairwise sequence alignment of full-length and predicted FH isoform based on experimentally verified transcript detected in RT-PCR.

**Supplementary Fig. 4:**
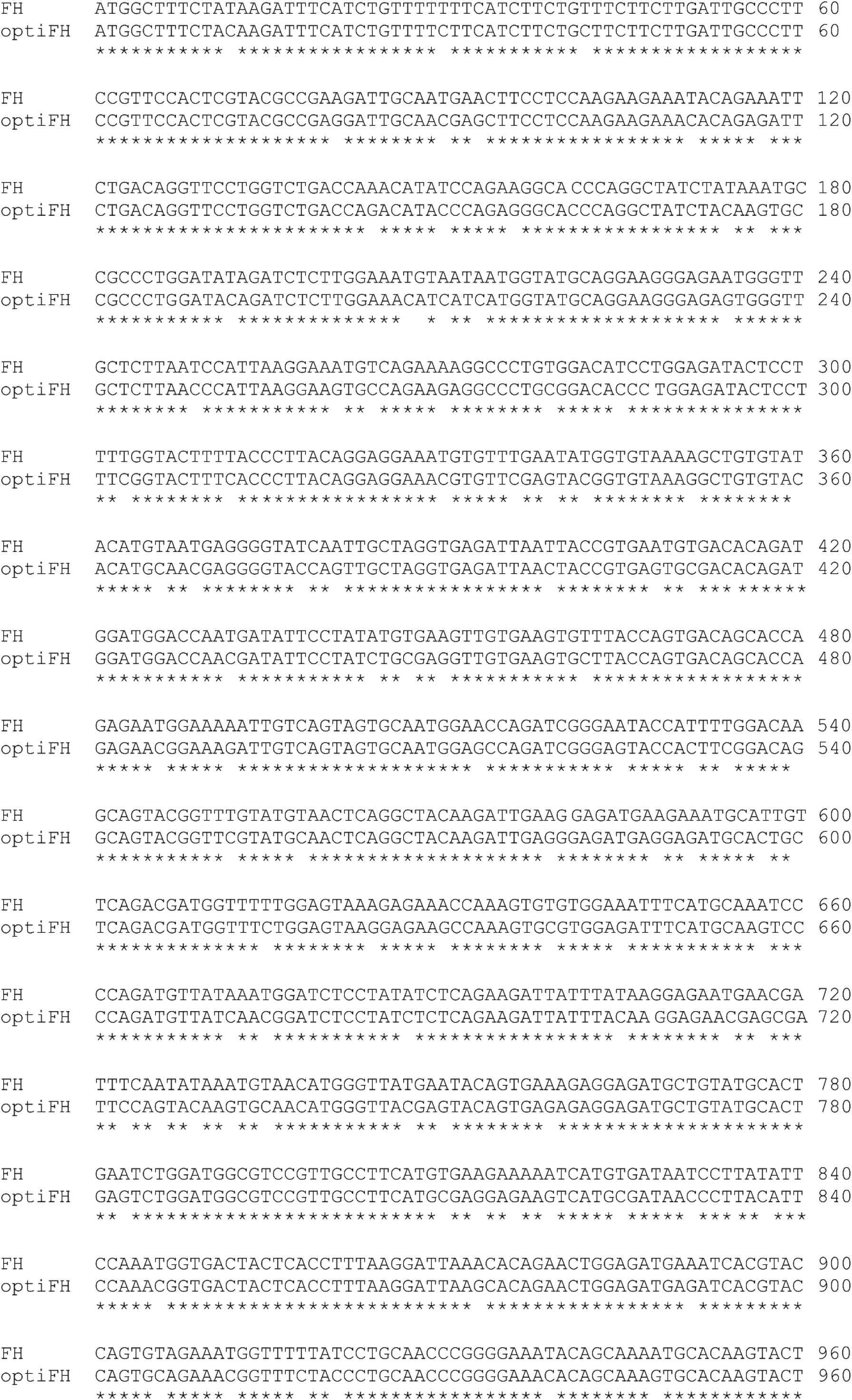
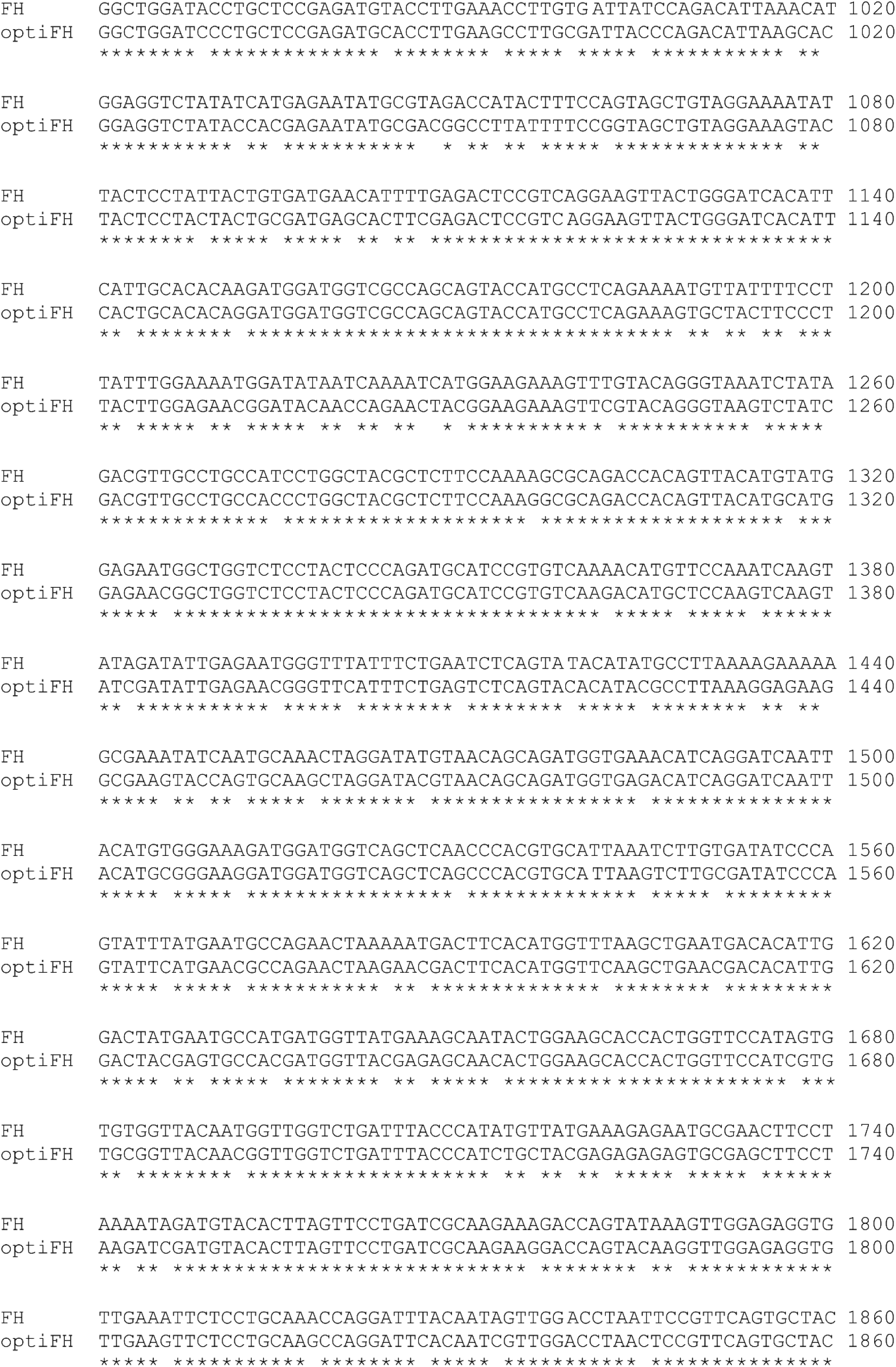
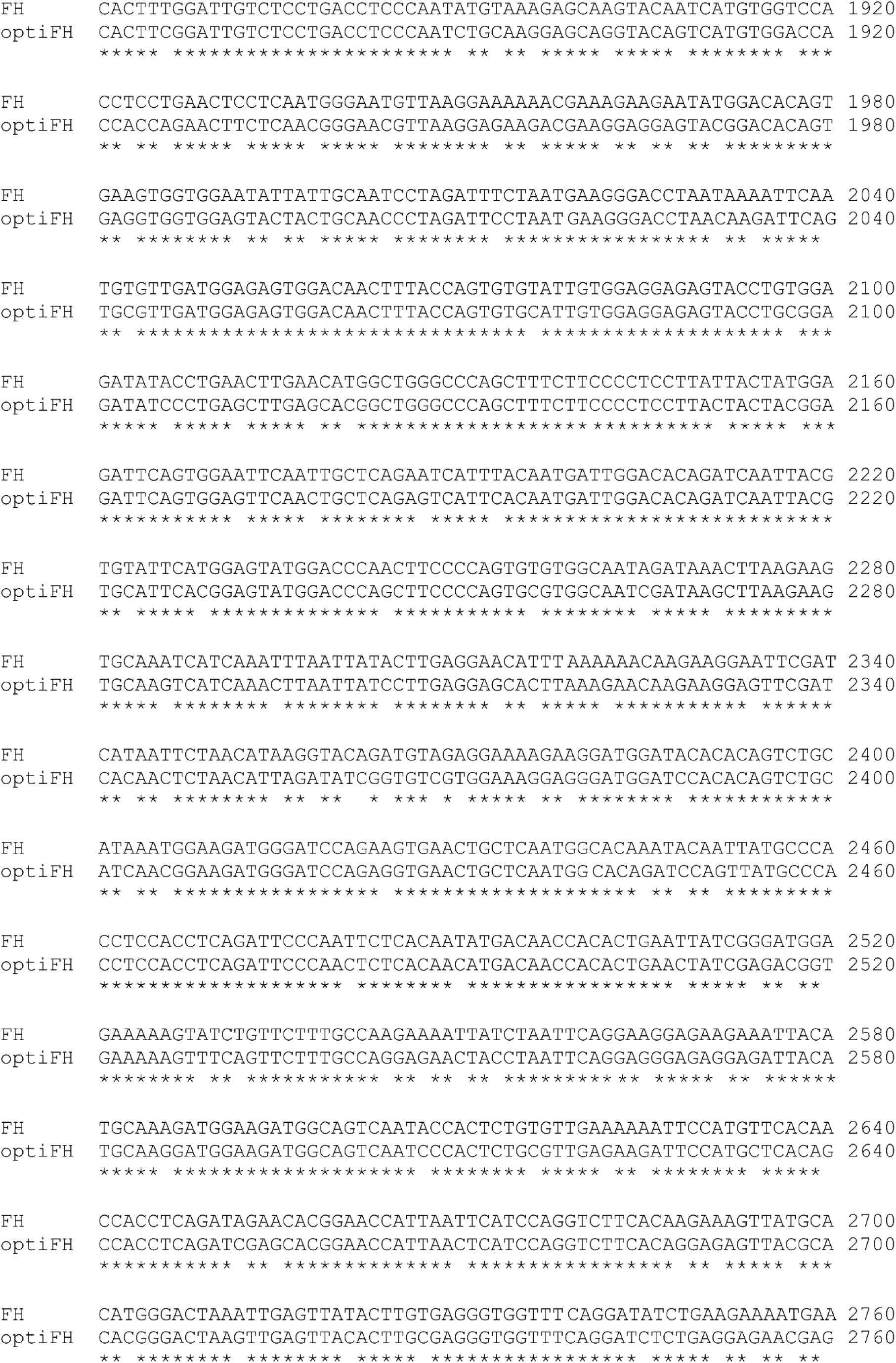
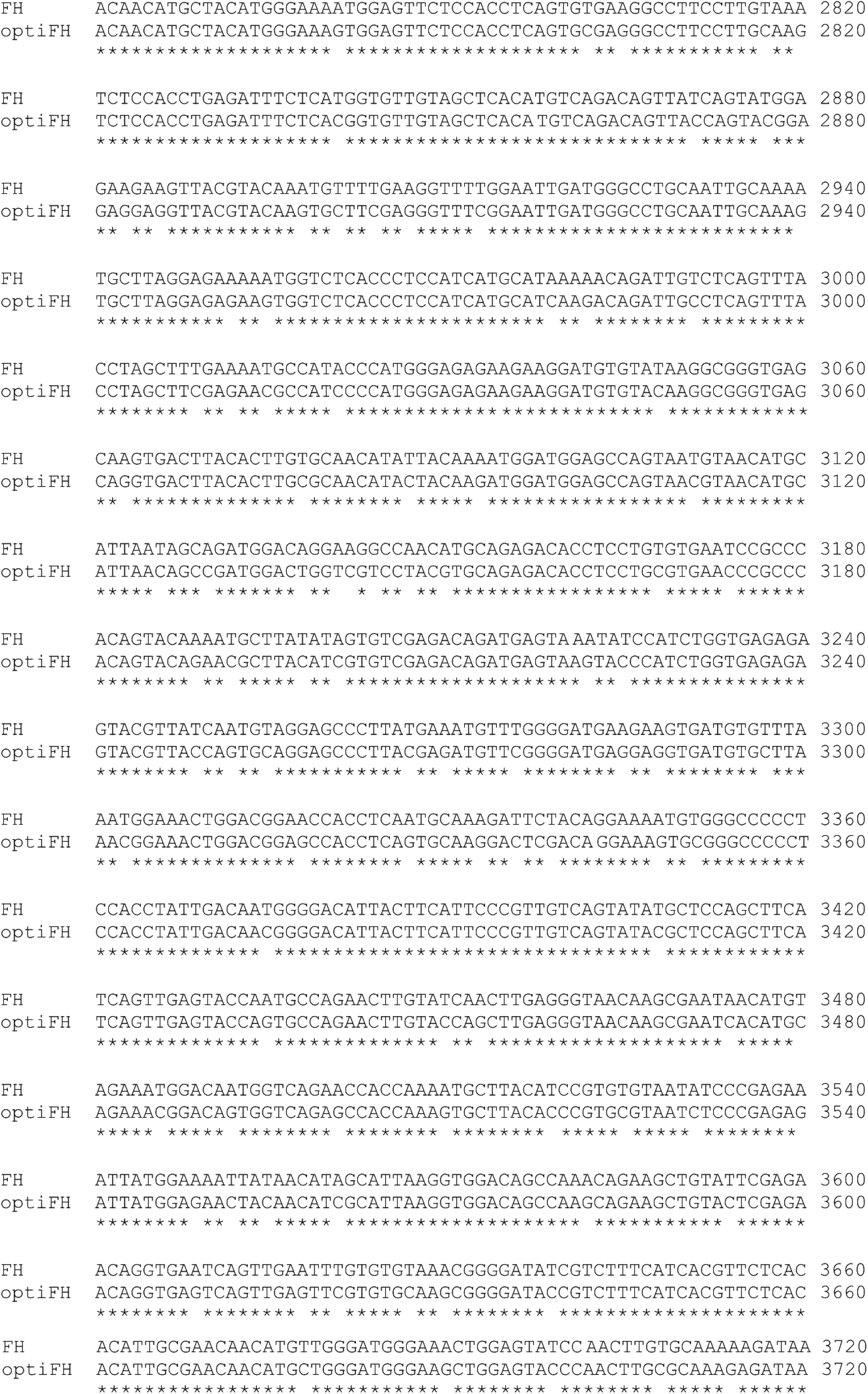
Pairwise sequence alignment of FH and optiFH.

**Supplementary Fig.5:**
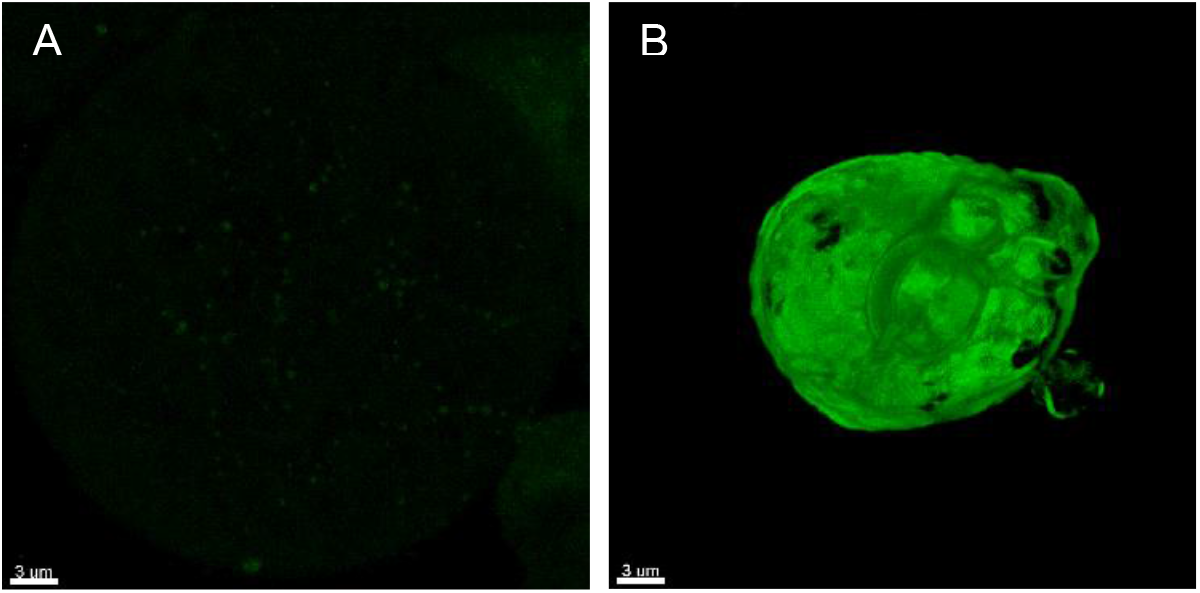
3D rendered confocal Z-stacks of moss protoplasts transfected with FH-Citrine (A) and optiFH-Citrine (B). Scale bar is 3 µm.

**Supplementary Fig. 6:**
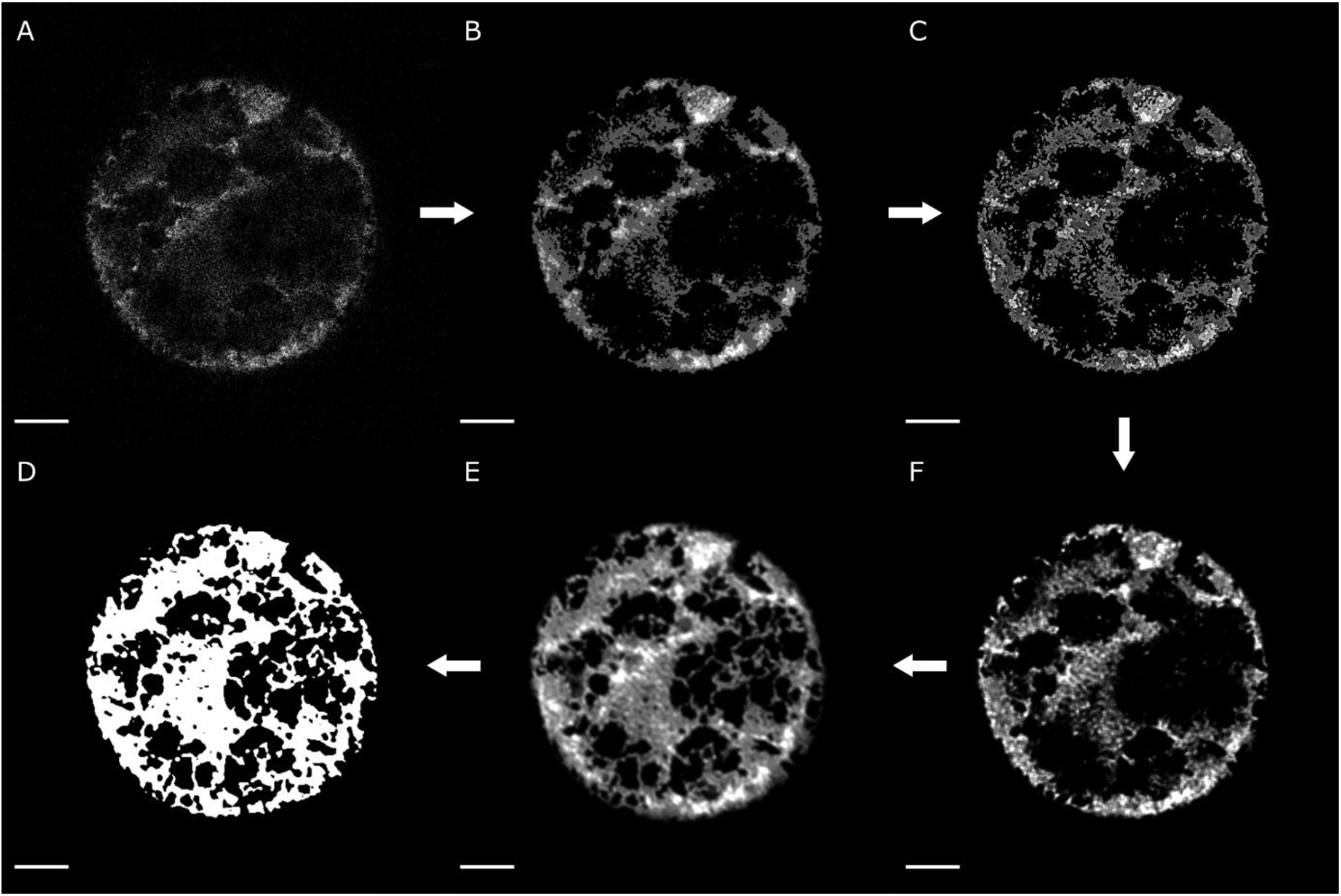
The workflow of the image processing steps demonstrated with an exemplary single slice from a FIX-Citrine cell. The original, raw image from the microscope (A) was first processed with a median filter (B). The resulting image was subjected to an unsharp-mask operation (C). Afterwards, a Richardson-Lucy restoration algorithm was implemented (F). The pre-processing was completed with a local intensity equalization operation (E). The final version of the image was then thresholded using a local adaptive Otsu thresholding method, leading to the binary masks (D) that were subsequently used for selection of the voxels for quantification. Scale bar is 5 µm.

**Supplementary Table 1:**
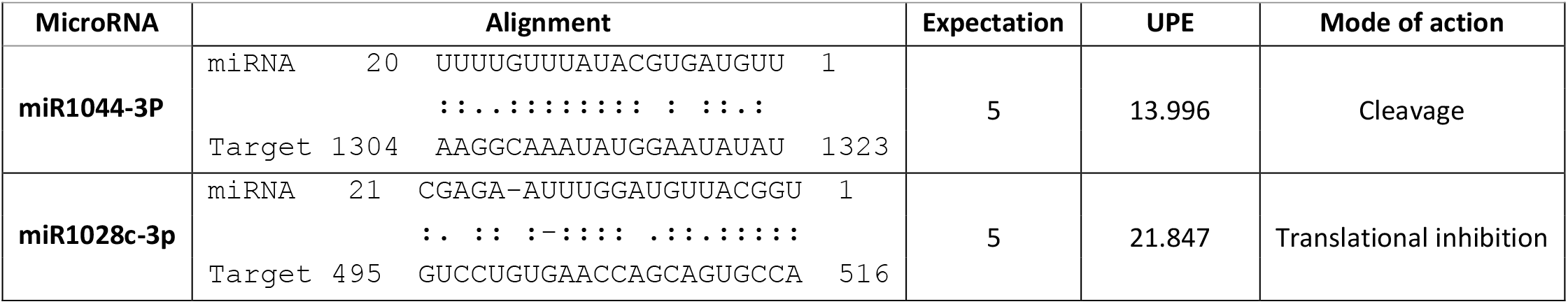
Moss microRNAs targeting human FIX CDS. Analysis was performed using psRNAtarget (http://plantgrn.noble.org/v1_psRNATarget/). The miRNAs as well as their binding sites, expectation score, maximum energy to unpair the target site (UPE) and mode of action of miRNAs are shown.

**Supplementary Table 2:** The donor splice sites for all 87,533 annotated transcripts corresponding to the 32,926 protein-encoding genes of the current *P. patens* genome release v3.3.

| *Motif* | *Count* | *Percentage (%)* | *Motif* | *Count* | *Percentage (%)* | *Motif* | *Count* | *Percentage (%)* |
| --- | --- | --- | --- | --- | --- | --- | --- | --- |
| CAGGT | 113,285 | 22.5460 | TAAGT | 1,610 | 0.3204 | AGAGC | 44 | 0.0088 |
| AAGGT | 84,769 | 16.8708 | AGTGT | 1,530 | 0.3045 | GCTGC | 44 | 0.0088 |
| GAGGT | 57,630 | 11.4695 | TGAGT | 1,492 | 0.2969 | ATCAT | 41 | 0.0082 |
| CTGGT | 20,698 | 4.1193 | AAGGC | 1,487 | 0.2959 | CGGGC | 41 | 0.0082 |
| ATGGT | 18,574 | 3.6966 | CTCGT | 1,385 | 0.2756 | GATGC | 41 | 0.0082 |
| TTGGT | 15,064 | 2.9980 | GGAGT | 1,330 | 0.2647 | TTTGC | 40 | 0.0080 |
| TAGGT | 12,835 | 2.5544 | ATCGT | 1,273 | 0.2534 | TCGGC | 39 | 0.0078 |
| CAAGT | 12,206 | 2.4292 | GACGT | 1,273 | 0.2534 | GCGGC | 36 | 0.0072 |
| GTGGT | 9,443 | 1.8793 | GTTGT | 1,259 | 0.2506 | GGGGC | 36 | 0.0072 |
| ACGGT | 9,246 | 1.8401 | TTCGT | 985 | 0.1960 | GGAGC | 35 | 0.0070 |
| TGGGT | 9,034 | 1.7980 | GAGGC | 981 | 0.1952 | CATGC | 34 | 0.0068 |
| AGGGT | 8,314 | 1.6547 | ACCGT | 914 | 0.1819 | TCAGC | 33 | 0.0066 |
| TCGGT | 7,588 | 1.5102 | TACGT | 908 | 0.1807 | CCGGC | 32 | 0.0064 |
| CGGGT | 6,757 | 1.3448 | CTAGT | 879 | 0.1749 | TTCGC | 32 | 0.0064 |
| AAAGT | 6,677 | 1.3289 | CGTGT | 870 | 0.1731 | ATTGC | 31 | 0.0062 |
| CCGGT | 6,223 | 1.2385 | ATAGT | 856 | 0.1704 | CTTGC | 31 | 0.0062 |
| GCGGT | 6,059 | 1.2059 | CCCGT | 804 | 0.1600 | GGTGC | 30 | 0.0060 |
| AATGT | 5,632 | 1.1209 | TCCGT | 742 | 0.1477 | TCTGC | 30 | 0.0060 |
| CATGT | 5,495 | 1.0936 | TGCGT | 673 | 0.1339 | AGTGC | 28 | 0.0056 |
| GGGGT | 5,371 | 1.0689 | AGCGT | 665 | 0.1323 | TGTGC | 26 | 0.0052 |
| GAAGT | 4,718 | 0.9390 | GCCGT | 516 | 0.1027 | CTCGC | 23 | 0.0046 |
| GATGT | 4,086 | 0.8132 | TTAGT | 506 | 0.1007 | TGCGC | 23 | 0.0046 |
| CTTGT | 3,916 | 0.7794 | GTCGT | 406 | 0.0808 | ACTGC | 22 | 0.0044 |
| ATTGT | 3,638 | 0.7240 | CGCGT | 396 | 0.0788 | GAAAT | 22 | 0.0044 |
| AGAGT | 2,987 | 0.5945 | GGTGT | 336 | 0.0669 | CCTGC | 21 | 0.0042 |
| CCTGT | 2,979 | 0.5929 | GTAGT | 300 | 0.0597 | GGCGC | 19 | 0.0038 |
| ACTGT | 2,904 | 0.5780 | GGCGT | 244 | 0.0486 | TACGC | 19 | 0.0038 |
| CAGGC | 2,860 | 0.5692 | CTGGC | 221 | 0.0440 | ACAGC | 18 | 0.0036 |
| GCTGT | 2,830 | 0.5632 | ATGGC | 199 | 0.0396 | GTTGC | 18 | 0.0036 |
| TCTGT | 2,772 | 0.5517 | TAGGC | 181 | 0.0360 | ATCGC | 17 | 0.0034 |
| TTTGT | 2,553 | 0.5081 | TTGGC | 89 | 0.0177 | TTAGC | 17 | 0.0034 |
| ACAGT | 2,426 | 0.4828 | CAAGC | 65 | 0.0129 | TGAGC | 16 | 0.0032 |
| CCAGT | 2,286 | 0.4550 | AAAGC | 55 | 0.0109 | AACGC | 15 | 0.0030 |
| CACGT | 2,273 | 0.4524 | GAAGC | 55 | 0.0109 | CCAGC | 15 | 0.0030 |
| AACGT | 1,965 | 0.3911 | TGGGC | 54 | 0.0107 | GACGC | 15 | 0.0030 |
| GCAGT | 1,948 | 0.3877 | AATGC | 52 | 0.0103 | GTCGC | 15 | 0.0030 |
| TATGT | 1,894 | 0.3769 | GTGGC | 52 | 0.0103 | GTCAT | 14 | 0.0028 |
| TCAGT | 1,882 | 0.3746 | ACGGC | 47 | 0.0094 | TATGC | 14 | 0.0028 |
| CGAGT | 1,876 | 0.3734 | AGGGC | 46 | 0.0092 | GTAGC | 13 | 0.0026 |
| TGTGT | 1,656 | 0.3296 | GCAGC | 46 | 0.0092 | TCCGC | 13 | 0.0026 |

| *Motif* | *Count* | *Percentage (%)* | *Motif* | *Count* | *Percentage (%)* |
| --- | --- | --- | --- | --- | --- |
| AAGAT | 12 | 0.0024 | TGCAT | 4 | 0.0008 |
| AGCGC | 12 | 0.0024 | AGAAT | 3 | 0.0006 |
| ATAGC | 12 | 0.0024 | ATGAT | 3 | 0.0006 |
| CGTGC | 12 | 0.0024 | GCTAT | 3 | 0.0006 |
| CTAGC | 12 | 0.0024 | TAATG | 3 | 0.0006 |
| CGAGC | 11 | 0.0022 | TTGAT | 3 | 0.0006 |
| AAAAT | 10 | 0.0020 | ATAAT | 2 | 0.0004 |
| CAAAT | 10 | 0.0020 | CAACT | 2 | 0.0004 |
| CACGC | 10 | 0.0020 | CCAAT | 2 | 0.0004 |
| CCCGC | 10 | 0.0020 | GACTA | 2 | 0.0004 |
| GCCGC | 10 | 0.0020 | GATAT | 2 | 0.0004 |
| ACCAT | 9 | 0.0018 | TAATA | 2 | 0.0004 |
| TAAGC | 9 | 0.0018 | TGAAA | 2 | 0.0004 |
| TCAAT | 9 | 0.0018 | TGGAT | 2 | 0.0004 |
| ATTAT | 8 | 0.0016 | CTGAT | 1 | 0.0002 |
| GCAAT | 8 | 0.0016 | CTGTG | 1 | 0.0002 |
| TCTAT | 8 | 0.0016 | CTTAT | 1 | 0.0002 |
| CATAT | 6 | 0.0012 | GGTAA | 1 | 0.0002 |
| TAGAT | 6 | 0.0012 | TAAAT | 1 | 0.0002 |
| TATAT | 6 | 0.0012 | TAACA | 1 | 0.0002 |
| ACCGC | 5 | 0.0010 | TAAGA | 1 | 0.0002 |
| CAGAT | 5 | 0.0010 | TAGAG | 1 | 0.0002 |
| CGCGC | 5 | 0.0010 | TGATG | 1 | 0.0002 |
| TCGAT | 5 | 0.0010 | TGCTT | 1 | 0.0002 |
| TTCAT | 5 | 0.0010 | TTGTC | 1 | 0.0002 |
| TTTAT | 5 | 0.0010 | TTTCG | 1 | 0.0002 |
| TAAGG | 4 | 0.0008 | TTTTA | 1 | 0.0002 |
| TGAGG | 4 | 0.0008 | TTTTT | 1 | 0.0002 |

**Supplementary Table 3:** The acceptor splice sites for all 87,533 annotated transcripts corresponding to the 32,926 protein-encoding genes of the current *P. patens* genome release v3.3.

| *Motif* | *Count* | *Percentage (%)* | *Motif* | *Count* | *Percentage (%)* |
| --- | --- | --- | --- | --- | --- |
| **CAGGT** | 90,834 | 18.0778 | **AAGTG** | 669 | 0.1331 |
| **CAGGA** | 54,132 | 10.7734 | **TAGCG** | 609 | 0.1212 |
| **CAGAT** | 45,474 | 9.0503 | **AAGAG** | 591 | 0.1176 |
| **CAGGG** | 36,713 | 7.3066 | **AAGAC** | 547 | 0.1089 |
| **CAGGC** | 36,350 | 7.2344 | **AAGCA** | 440 | 0.0876 |
| **CAGAA** | 22,672 | 4.5122 | **GAGGT** | 365 | 0.0726 |
| **CAGAG** | 21,613 | 4.3014 | **AAGCC** | 315 | 0.0627 |
| **TAGGT** | 19,555 | 3.8918 | **AAGTA** | 285 | 0.0567 |
| **CAGTT** | 19,239 | 3.8290 | **AAGTC** | 273 | 0.0543 |
| **CAGCT** | 18,854 | 3.7523 | **AAGCG** | 238 | 0.0474 |
| **CAGTG** | 15,897 | 3.1638 | **GAGGA** | 185 | 0.0368 |
| **CAGAC** | 14,770 | 2.9395 | **GAGAT** | 182 | 0.0362 |
| **CAGCA** | 11,526 | 2.2939 | **GAGGG** | 168 | 0.0334 |
| **TAGGA** | 9,029 | 1.7970 | **GAGGC** | 160 | 0.0318 |
| **CAGCC** | 8,266 | 1.6451 | **GAGAA** | 126 | 0.0251 |
| **CAGTC** | 7,702 | 1.5329 | **GAGTG** | 108 | 0.0215 |
| **TAGGG** | 7,378 | 1.4684 | **GAGAG** | 100 | 0.0199 |
| **TAGAT** | 7,164 | 1.4258 | **GAGCT** | 80 | 0.0159 |
| **CAGTA** | 6,745 | 1.3424 | **GAGTT** | 77 | 0.0153 |
| **TAGGC** | 6,040 | 1.2021 | **GAGAC** | 69 | 0.0137 |
| **CAGCG** | 6,004 | 1.1949 | **GAGCA** | 67 | 0.0133 |
| **AAGGT** | 4,063 | 0.8086 | **CACAT** | 51 | 0.0102 |
| **TAGAG** | 2,667 | 0.5308 | **GAGTC** | 48 | 0.0096 |
| **TAGAA** | 2,624 | 0.5222 | **GAGTA** | 46 | 0.0092 |
| **TAGTT** | 2,554 | 0.5083 | **GAGCG** | 39 | 0.0078 |
| **TAGTG** | 2,225 | 0.4428 | **GAGCC** | 37 | 0.0074 |
| **TAGCT** | 2,183 | 0.4345 | **CACGC** | 26 | 0.0052 |
| **TAGAC** | 2,119 | 0.4217 | **CACAA** | 16 | 0.0032 |
| **AAGGA** | 1,792 | 0.3566 | **TACCT** | 14 | 0.0028 |
| **AAGGC** | 1,483 | 0.2951 | **CACAC** | 11 | 0.0022 |
| **TAGCA** | 1,480 | 0.2946 | **CACGG** | 10 | 0.0020 |
| **AAGAT** | 1,342 | 0.2671 | **TACAC** | 10 | 0.0020 |
| **AAGGG** | 1,107 | 0.2203 | **CACGT** | 8 | 0.0016 |
| **TAGTC** | 967 | 0.1925 | **TACAT** | 8 | 0.0016 |
| **TAGCC** | 916 | 0.1823 | **CACAG** | 6 | 0.0012 |
| **AAGAA** | 784 | 0.1560 | **TACAA** | 6 | 0.0012 |
| **TAGTA** | 757 | 0.1507 | **TACGA** | 6 | 0.0012 |
| **AAGTT** | 685 | 0.1363 | **TACGC** | 6 | 0.0012 |
| **AAGCT** | 671 | 0.1335 | **AACGT** | 5 | 0.0010 |
| AGAAT | 5 | 0.0010 | ACGAT | 1 | 0.0002 |
| CACCT | 4 | 0.0008 | AGCAT | 1 | 0.0002 |
| AACAA | 3 | 0.0006 | ATAAT | 1 | 0.0002 |
| AGGAT | 3 | 0.0006 | CATCT | 1 | 0.0002 |
| CACCA | 3 | 0.0006 | CTAAT | 1 | 0.0002 |
| CACTA | 3 | 0.0006 | CTGGC | 1 | 0.0002 |
| TACGT | 3 | 0.0006 | GATAT | 1 | 0.0002 |
| TACTC | 3 | 0.0006 | GCTAT | 1 | 0.0002 |
| AACAG | 2 | 0.0004 | GCTTC | 1 | 0.0002 |
| AAC TT | 2 | 0.0004 | GGAAG | 1 | 0.0002 |
| ACTAA | 2 | 0.0004 | GGTAT | 1 | 0.0002 |
| CACTG | 2 | 0.0004 | GTGAT | 1 | 0.0002 |
| CACTT | 2 | 0.0004 | GTTAT | 1 | 0.0002 |
| GAAAT | 2 | 0.0004 | NAGGA | 1 | 0.0002 |
| GAACT | 2 | 0.0004 | NAGGG | 1 | 0.0002 |
| GACAA | 2 | 0.0004 | NAGTG | 1 | 0.0002 |
| GACAT | 2 | 0.0004 | TACCG | 1 | 0.0002 |
| GCAAT | 2 | 0.0004 | TACTA | 1 | 0.0002 |
| GCCAT | 2 | 0.0004 | TACTG | 1 | 0.0002 |
| GGCAT | 2 | 0.0004 | TGCGT | 1 | 0.0002 |
| GGGAT | 2 | 0.0004 | TGTGA | 1 | 0.0002 |
| AACAT | 1 | 0.0002 | TTATT | 1 | 0.0002 |
| ACCAT | 1 | 0.0002 | TTTAC | 1 | 0.0002 |

